# Mutant lamins cause nuclear envelope rupture and DNA damage in skeletal muscle cells

**DOI:** 10.1101/364778

**Authors:** Ashley J. Earle, Tyler J. Kirby, Gregory R. Fedorchak, Philipp Isermann, Jineet Patel, Sushruta Iruvanti, Steven A. Moore, Gisèle Bonne, Lori L. Wallrath, Jan Lammerding

## Abstract

Mutations in the human *LMNA* gene, which encodes the nuclear envelope (NE) proteins lamins A and C, cause autosomal dominant Emery-Dreifuss muscular dystrophy, congenital muscular dystrophy, limb-girdle muscular dystrophy, and other diseases collectively known as laminopathies. The molecular mechanisms responsible for these diseases remain incompletely understood, but the muscle-specific defects suggest that mutations may render nuclei more susceptible to mechanical stress. Using three mouse models of muscle laminopathies, we found that *Lmna* mutations caused extensive NE abnormalities, consisting of chromatin protrusions into the cytoplasm and transient rupture of the NE in skeletal muscle cells. NE damage was associated with DNA damage, activation of DNA damage response pathways, and reduced viability. Intriguingly, NE damage resulted from nuclear migration in maturing skeletal muscle cells, rather than actomyosin contractility. NE damage and DNA damage was reduced by either depletion of kinesin-1 or disruption of the Linker of Nucleoskeleton and Cytoskeleton (LINC) complex. LINC complex disruption rescued myofiber function and viability in *Lmna* mutant myofibers, indicating that the myofiber dysfunction is the result of mechanically induced NE damage. The extent of NE damage and DNA damage in *Lmna* mouse models correlated with the disease onset and severity *in vivo*. Moreover, inducing DNA damage in wild-type muscle cells was sufficient to phenocopy the reduced cell viability of lamin A/C-deficient muscle cells, suggesting a causative role of DNA damage in disease pathogenesis. Corroborating the mouse model data, muscle biopsies from patients with *LMNA* muscular dystrophy revealed significant DNA damage compared to age-matched controls, particularly in severe cases of the disease. Taken together, these findings point to a new and important role of DNA damage as a pathogenic contributor for *LMNA* skeletal muscle diseases.

## INTRODUCTION

Lamins A and C, together with the B-type lamins, are the major components of the nuclear lamina, which line the inner nuclear membrane. Lamins A/C play important roles in providing structural support to the nucleus and connecting the nucleus to the cytoskeleton^2^. In addition, they participate in transcriptional regulation, genome organization, and DNA damage and repair^2–5^. The majority of the over 450 *LMNA* mutations identified to date are responsible for autosomal dominant Emery-Dreifuss muscular dystrophy (AD-EDMD)^6^, characterized by slowly progressive skeletal muscle wasting, contractures of the elbow, neck, and Achilles tendons, a rigid spine, abnormal heart rhythms, heart block, and cardiomyopathy^7^. Other *LMNA* mutations cause congenital muscular dystrophy (*LMNA*-CMD), a particularly severe form of muscular dystrophy with onset in early childhood^8^, and limb girdle muscular dystrophy, which affects proximal muscles of the hips and shoulders^9^. It remains unclear how *LMNA* mutations result in muscle-specific defects, and the incomplete understanding of the disease pathogenesis presents a major hurdle in the development of effective treatment approaches.

One potential explanation, the ‘mechanical stress’ hypothesis, states that *LMNA* mutations linked to muscular phenotypes result in structurally impaired nuclei that become damaged in mechanically active tissues, such as cardiac and skeletal muscle^5^. This hypothesis is supported by findings of decreased nuclear stiffness in fibroblasts expressing *LMNA* mutations linked to striated muscle laminopathies, impaired assembly of mutant lamins *in vitro*, and anecdotal reports of NE damage in skeletal and cardiac muscle cells of individuals with AD-EDMD and *LMNA*-related dilated cardiomyopathies^2, 5, 10, 11^. However, striated muscle laminopathies involve progressive muscle wasting, and analysis of fixed tissues and cells provides only a snapshot of the development of disease. Thus, the functional relevance of NE damage, particularly whether the NE damage is a cause or consequence of muscle dysfunction, remains unclear.

To better understand the mechanistic link between NE damage and muscle dysfunction, we employed three laminopathy mouse models with varying degrees of disease severity, along with a recently developed long-term *in vitro* muscle differentiation platform^12^. Using high resolution time lapse microscopy, we tracked myonuclear shape and integrity over time and recorded structural and functional changes during the differentiation of primary mouse myoblasts into multinucleated myotubes and subsequent formation of mature, contractile myofibers (Fig. 1a, b). Combining these models with novel, fluorescent nuclear damage reporters^13^, both *in vitro* and *in vivo*, we obtained unprecedented systematic and detailed temporal and mechanistic information of disease progression in mouse models of laminopathies, and corroborated key findings in skeletal muscle biopsies from humans with *LMNA* muscular dystrophy.

**Figure 1.**
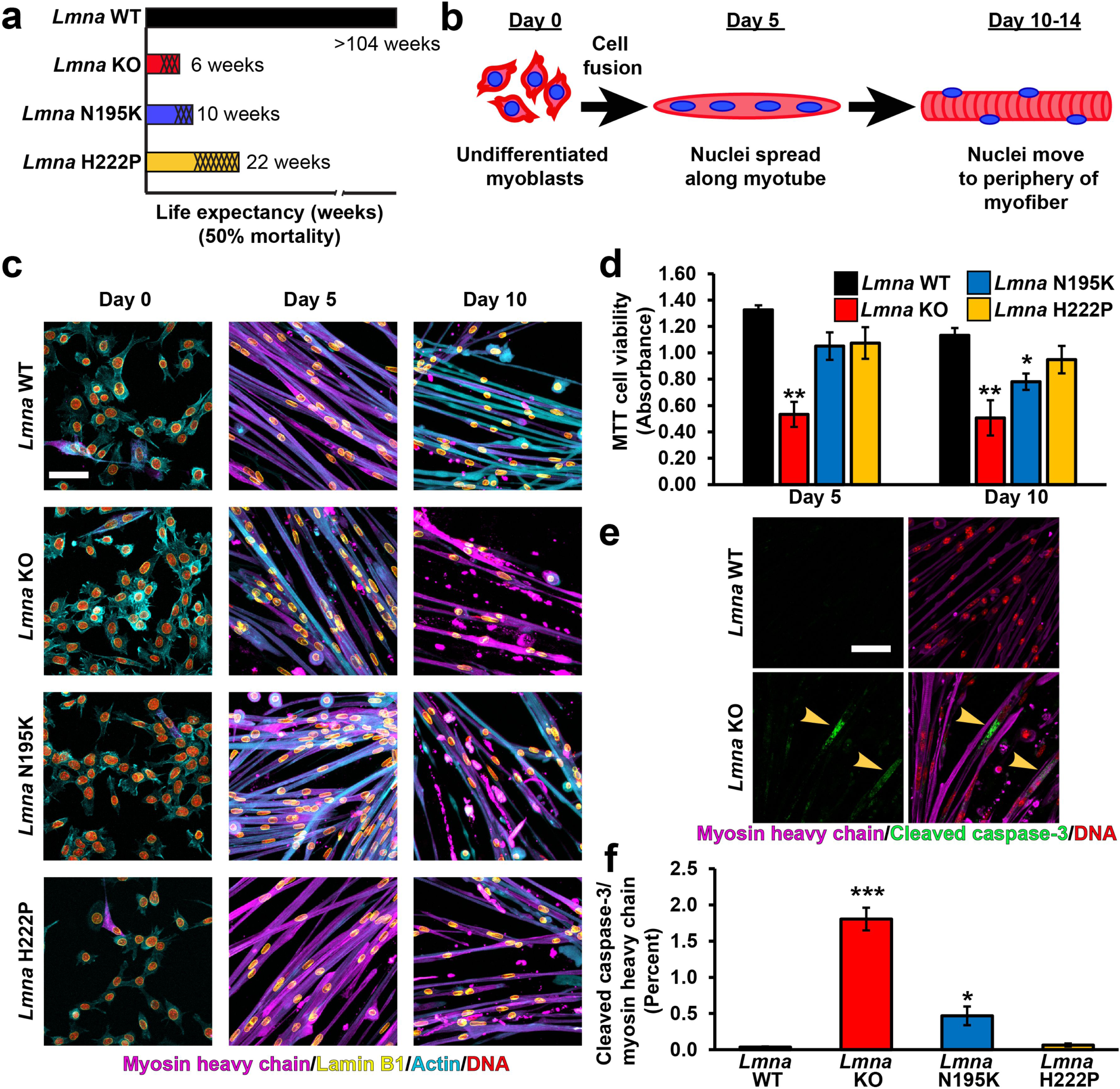
***In vitro* differentiated primary myoblasts from *Lmna* KO, *Lmna* N195K, and *Lmna* H222P mice recapitulates disease severity**. (**a**) Graphical representation of the three *Lmna* mutant models used in the study, indicating the published 50% mortality rates of *Lmna* KO, *Lmna* N195K, and *Lmna* H222P mice, as well as wild-type (*Lmna* WT) controls. Shading represents the onset of disease symptoms in the mouse models. (**b**) Schematic for the stages of differentiation from primary myoblasts into mature myofibers in the *in vitro* system. (**c**) Representative images of *Lmna* WT, *Lmna* KO, *Lmna* N195K and *Lmna* H222P primary skeletal muscle cells at days 0, 5 and 10 of differentiation. Scale bar: 100 µm. (**d**) Quantification of cell viability using MTT assay at days 5, 10 of differentiation. *n* = 3-6 independent cell lines for each genotype. **, *p* < 0.01 vs. *Lmna* WT; *, *p* < 0.05 vs. *Lmna* WT. (**e**) Representative image of cleaved caspase-3 immunofluorescence in *Lmna* WT and *Lmna* KO myofibers at day 10 of differentiation. Scale bar: 20 µm (**f**) Quantification of cleaved caspase-3 relative to myosin heavy chain immunofluorescence area in *Lmna* WT, *Lmna* KO, *Lmna* N195K and *Lmna* H222P myofibers after 10 days of differentiation ***, *p* < 0.001 vs. *Lmna* WT; *, *p* < 0.05 vs. *Lmna* WT. *n* = 3 independent cell lines for each genotype.

We found that myonuclei in *Lmna* mutant muscle cells exhibited progressive NE damage *in vitro* and *in vivo*, including extensive chromatin protrusions into the cytoplasm and transient NE rupture. Intriguingly, NE rupture was associated with progressive DNA damage and DNA damage response activation. Inducing DNA damage in wild-type muscle cells was sufficient to provoke defects in cell viability similar to those observed in the lamin A/C-deficient cells. In addition, preventing NE rupture by LINC complex disruption was sufficient to reduce DNA damage and rescue myofiber viability and contractility in lamin A/C-deficient cells. Together, our findings indicate a causative role of NE rupture and DNA damage in progressive muscle decline and provide a novel explanation for how lamin mutations lead to muscle weakness and wasting in AD-EDMD.

## RESULTS

### *Lmna* mutations cause progressive decline in myofiber health *in vitro* and *in vivo*

To examine the effect of *Lmna* mutations on nuclear mechanics and muscle function *in vitro*, we isolated myoblasts from three established mouse models of striated muscle laminopathies, representing a spectrum of muscle wasting and disease severity (**Fig. 1a,** Suppl. Fig. S1): Lamin A/C-deficient (*Lmna*^–/–^) mice^14^, subsequently referred to as lamin A/C knock-out mice (*Lmna* KO); knock-in mice carrying the *Lmna*^N195K/N195K^ mutation (*Lmna* N195K)^15^; knock-in mice carrying the *Lmna*^H222P/H222P^ mutation (*Lmna* H222P)^16^; and wild-type littermates. While the *Lmna* N195K mice were originally described as a model for dilated cardiomyopathy^15^, in the C57BL/6 background used in our studies, the mice developed pronounced skeletal muscular dystrophy in addition to cardiac defects (Suppl. Fig. S1). For *in vitro* studies, we utilized a recently developed, three-dimensional culture protocol to differentiate primary myoblasts into mature, contractile myofibers over the course of ten days (**Fig. 1b**)^17, 18^. The resulting myofibers display the highly organized sarcomeric structure and evenly spaced peripheral myonuclear positioning characteristic of mature skeletal muscle fibers *in vivo* (Suppl. Fig. S2).

All myoblasts, including the *Lmna* KO, *Lmna* N195K and *Lmna* H222P mutant cells, successfully completed myoblast fusion, differentiated into myotubes, and matured into myofibers (**Fig. 1c**), consistent with previous studies on differentiation of *Lmna* KO myoblasts^19, 20^. Wild-type myofibers remained healthy and highly contractile up to ten days of differentiation

(Suppl. Movie 1). In contrast, the *Lmna* KO myofibers showed a decline in cell contractility, viability, and number of myonuclei, starting at day five of differentiation (**Fig. 1c, d**; Suppl. Fig. S3, Suppl. Movie 2). The *Lmna* N195K myofibers showed a similar, albeit slightly delayed decline in cell viability, contractility, and number of myonuclei by day ten of differentiation (**Fig. 1d**, Suppl. Fig S3, Suppl. Movie 3). The reduction in cell viability at day ten in the *Lmna* KO and *Lmna* N195K myofibers was associated with an increase in activated caspase-3 (**Fig. 1e,f**), indicating that reduced viability was due at least in part to cell-intrinsic apoptosis. Unlike the *Lmna* KO and *Lmna* N195K models, the *Lmna* H222P myofibers exhibited no significant decrease in viability, contractility, or number of myonuclei (**Fig. 1d**, Suppl. Fig S3, Suppl. Movie 4) within ten days of differentiation. Taken together, the long-term differentiation assays revealed a striking correlation among the defects observed *in vitro*, including loss of muscle cell viability and presence of apoptotic markers, with the severity of the disease in the corresponding mouse models (**Fig 1a**, Suppl. Fig. S1), suggesting that defects in the *in vitro* model may serve as prognostic markers for disease progression.

### *Lmna* mutant muscle cells have reduced nuclear stability that corresponds to disease severity

We hypothesized that the progressive deterioration of *Lmna* mutant myofibers results from damage to mechanically weakened myonuclei exposed to cytoskeletal forces. To test this hypothesis, we measured the nuclear deformability in primary myoblasts from the three laminopathy models using a novel high throughput microfluidics-based micropipette aspiration assay (**Fig. 2a**, Suppl. Movie 5)^21^. Nuclei from *Lmna* KO and *Lmna* N195K myoblasts were substantially more deformable than nuclei from wild-type controls (**Fig. 2b,c**). Intriguingly, myoblasts from *Lmna* H222P mice, which have a later disease onset and less severe muscle defects than the other two *Lmna* mutant models (**Fig. 1a;** Suppl. Fig. S1), had only a modest increase in nuclear deformability relative to wild-type controls (**Fig. 2b,c**; Suppl. Fig. S4a). Ectopic expression of lamin A significantly reduced the nuclear deformability defect in primary *Lmna* KO myoblasts (Suppl. Fig. S4b-d), confirming that the impaired nuclear stability was a direct consequence of altering the nuclear lamina. In addition, primary myoblasts from *Mdx* mice, which develop mild muscular dystrophy due to loss of the cell membrane-associated protein dystrophin, had nuclear deformation indistinguishable from wild-type controls (Suppl. Fig. S5), indicating that the defects in nuclear stability are specific to *Lmna* mutations and not muscular dystrophy in general.

**Figure 2.**
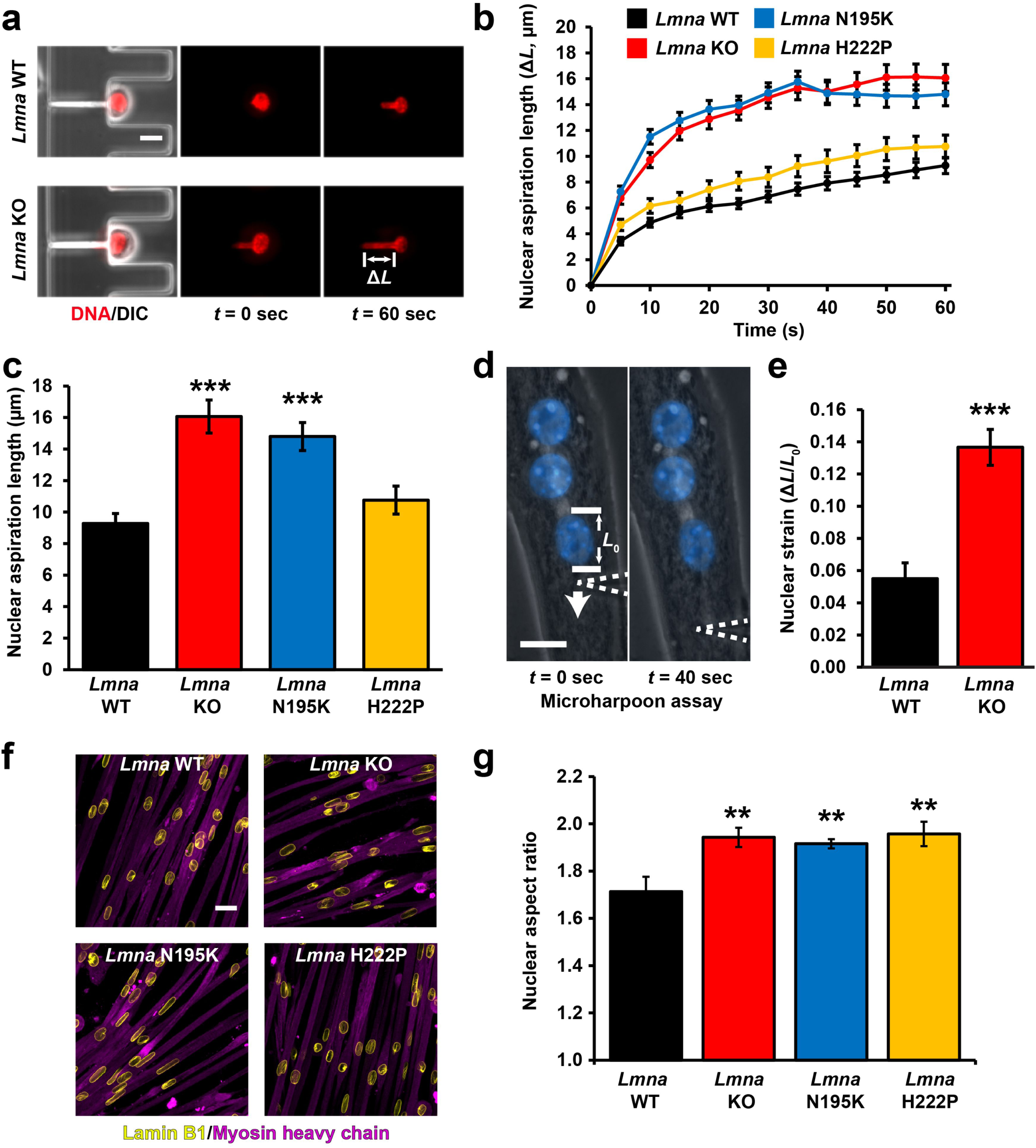
***Lmna* mutant muscle cells display defects in nuclear stability**. (**a**) Representative images of *Lmna* WT and *Lmna* KO nuclei deforming in a microfluidic micropipette aspiration device. Scale bar: 10 µm. (**b**) Measurement for nuclear deformation at 5 second intervals for *Lmna* WT, *Lmna* KO, *Lmna* N195K, and *Lmna* H222P myoblasts during 60 seconds of aspiration. (**c**) Quantification of the nuclear deformation after 60 seconds of aspiration. *n* = 41-67 nuclei per genotype from 3 independent experiments. ***, *p* < 0.001 *vs*. *Lmna* WT. (**d**) Microharpoon assay to measure nuclear deformability (Δ*L*/*L*_0_) in myofibers, showing representative images before and at the end of perinuclear cytoskeletal strain application with a microneedle (dashed line). Scale bar: 15 µm. (**e**) Quantification of nuclear strain induced by microharpoon assay in *Lmna* WT and *Lmna* KO myotubes at day 5 of differentiation. *n* = 19-22 nuclei per genotype from 3 independent experiments. ***, *p* < 0.001 *vs*. *Lmna* WT myotubes. (**f**) Representative image of nuclear morphology in *Lmna* WT, *Lmna* KO, *Lmna* N195K and *Lmna* H222P myotubes after 5 days of differentiation. Scale bar: 20 µm (**g**) Nuclear aspect ratio (length/width) in *Lmna* WT, *Lmna* KO, *Lmna* N195K and *Lmna* H222P myotubes after 5 days of differentiation. *n* = 3 - 4 independent cell lines per genotype with >100 nuclei counted per image. **, *p* < 0.01 *vs*. *Lmna* WT.

To assess whether the observed defects in nuclear stability also occur in more mature, multinucleated myofibers, we subjected *Lmna* KO and wild-type myofibers to a ‘microharpoon’ assay, in which precise strain is exerted on the perinuclear cytoskeleton, and the induced nuclear deformation and displacement are used to infer nuclear stability and nucleo-cytoskeletal coupling, respectively^22, 23^. *Lmna* KO myofibers had significantly more deformable nuclei than wild-type controls (**Fig. 2d,e**; Suppl. Movie 6), consistent with the micropipette aspiration results in the myoblasts. Furthermore, analysis of *Lmna* mutant and wild-type myofibers at day five of *in vitro* differentiation revealed that *Lmna* KO, *Lmna* N195K, and *Lmna* H222P myofibers had significantly elongated myonuclei compared to wild-type controls (**Fig. 2f,g**), consistent with decreased nuclear stability in the *Lmna* mutant cells and with previous reports of elongated nuclei in muscle biopsies from laminopathy patients^24^. Taken together, these findings suggest that myopathic *Lmna* mutations that cause severe muscle defects result in mechanically weaker myonuclei.

### *Lmna* mutant myonuclei display chromatin protrusions into the cytoplasm

Physical stress on the nucleus during external compression or confined migration can induce chromatin protrusions across the nuclear lamina into the cytoplasm, particularly in lamin-deficient cells^13, 25^. To test whether the mechanically weaker *Lmna* mutant myonuclei are prone to mechanically induced damage in muscle cells, we analyzed nuclear structure and morphology over the ten-day time course of differentiation and maturation of primary myoblasts from laminopathy mouse models and healthy controls. Despite their mechanically weaker nuclei (**Fig. 2**), the *Lmna* mutant myoblasts show no nuclear abnormalities prior to differentiation (**Fig. 3b**). Following the onset of differentiation, however, *Lmna* KO, *Lmna* N195K, and *Lmna* H222P myofibers exhibited striking chromatin protrusions that were absent in wild-type fibers. These protrusions extended beyond the (B-type) nuclear lamina up to tens of microns into the cytoplasm (**Fig. 3a,b**). The protrusions were enclosed by nuclear membranes, as indicated by the frequent presence of the nuclear membrane protein emerin, and occasional presence of nesprin-1 (Suppl. Fig. S6a); however, these NE proteins were often concentrated in punctae inside the protrusions and myonuclei. Other NE proteins, such as nuclear pore complex proteins, were largely absent from the protrusions (Suppl. Fig. S6b), suggesting an altered membrane composition in the chromatin protrusion, similar to what has been reported in analogous structures in migrating cancer cells^13, 26^.

**Figure 3.**
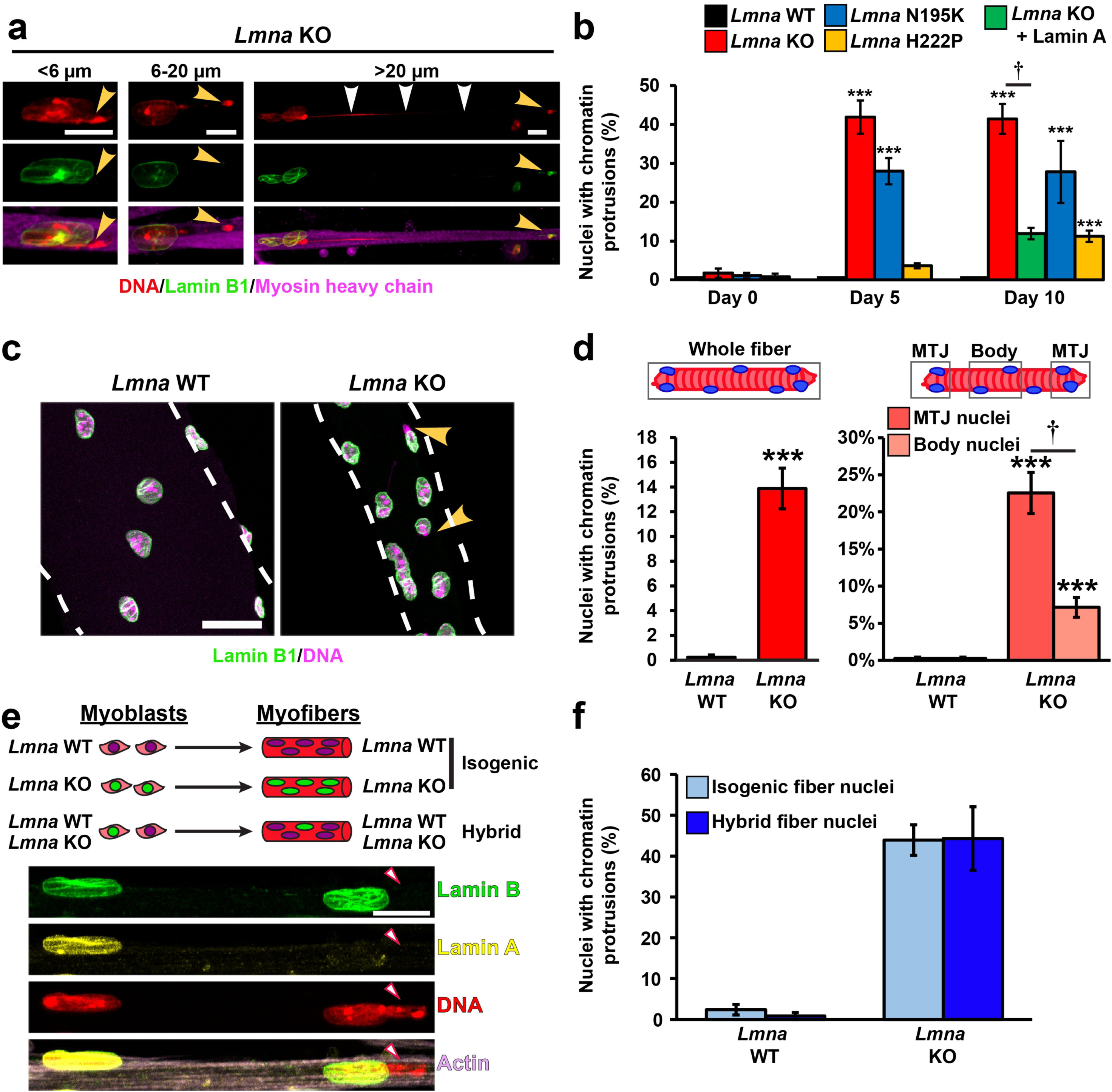
***Lmna* mutant myonuclei develop chromatin protrusions during differentiation.** (**a**) Representative images of chromatin protrusions observed in *Lmna* KO myofibers after 10 days of differentiation. Yellow arrowheads indict the end of the protrusion; the white arrowheads indicate a thin chromatin tether protruding from the nucleus. Scale bar: 10 µm. (**b**) Quantification of the percentage of myonuclei containing chromatin protrusion at days 5 and 10 of differentiation in *Lmna* WT, *Lmna* KO, *Lmna* KO + Lamin A, *Lmna* N195K and *Lmna* H222P cell lines. Data from *n* = 3 independent experiments with 62-73 nuclei per genotype. ***, *p* < 0.001 *vs*. *Lmna* WT cells. †, *p* < 0.01 *vs*. *Lmna* KO. (**c**) Representative images of isolated single muscle fibers from *Lmna* WT and *Lmna* KO mice labeled for lamin B1 (green) and DNA (magenta). Arrowheads indicate the presence of chromatin protrusions in *Lmna* KO muscle fiber. Scale bar: 20 µm. (**d**) Quantification of the percentage of myonuclei with chromatin protrusion in isolated muscle fibers from *Lmna* WT and *Lmna* KO mice. Left, data based on analysis of total muscle fiber. Right, analysis for nuclei located at the MTJ compared to those within the body of the fiber. *n* = 8-11 mice per genotype, with 5 single fibers imaged per animal. ***, *p* < 0.001 *vs*. *Lmna* WT. †, *p* < 0.01 *vs*. nuclei in the muscle body. (**e**) Top, schematic of the generation of hybrid myofibers containing nuclei from both *Lmna* WT and *Lmna* KO cell lines. Bottom, corresponding representative images. Final hybrid fibers contained ∼80% *Lmna* WT nuclei and 20% *Lmna* KO nuclei. Arrowheads denote *Lmna* KO nucleus with a chromatin protrusion residing within the same myofiber as a *Lmna* WT nucleus. (**f**) Quantification of the number of chromatin protrusions from *Lmna* WT and *Lmna* KO contained within isogenic myofibers (control) or hybrid myofibers containing 80% *Lmna* WT and 20% *Lmna* KO nuclei. *n* = 3 independent experiments, in which 91-163 nuclei were quantified per experiment.

The frequency of chromatin protrusion was highest in *Lmna* KO myofibers, followed by *Lmna* N195K and then *Lmna* H222P myofibers (**Fig. 3b**), correlating with the increased nuclear deformability *in vitro* (**Fig. 2**) and the disease severity *in vivo* (**Fig. 1a;** Suppl. Fig. S1). Intriguingly, while *Lmna* KO and *Lmna* N195K myofibers had extensive chromatin protrusions at day five of differentiation, the frequency of chromatin protrusions in the *Lmna* H222P cells was initially low, but increased from five to ten days of differentiation (**Fig. 3b**), matching the delayed disease onset and progressive phenotype of the *Lmna* H222P model *in vivo.* Ectopic expression of lamin A in *Lmna* KO myoblasts significantly reduced the occurrence of chromatin protrusions at ten days of differentiation (**Fig. 3b**), confirming that the protrusions were caused by loss of lamin A/C expression.

To confirm the results of the *in vitro* studies *in vivo*, we isolated single muscle fibers from the hindlimbs of *Lmna* KO and wild-type mice. Chromatin protrusions were not detectable in muscle fibers from wild-type mice (**Fig. 3c,d**). In contrast, nuclei in muscle fibers from *Lmna* KO mice had similar chromatin protrusions as observed in the *in vitro* differentiated myofibers (**Fig. 3c,d**). Interestingly, the prevalence of chromatin protrusions in the *Lmna* KO myonuclei strongly depended on the location within the muscle. Myonuclei at the myotendinous junctions (MTJ) had a higher frequency of chromatin protrusions than nuclei in the muscle fiber body (**Fig. 3d**), consistent with a previous report of nuclear abnormalities at the MTJ in *Lmna* KO mice^27^, and possibly reflecting increased mechanical stress at the MTJ.

## Nuclear damage is intrinsic to lamin A/C-deficient nuclei

To address whether the observed NE defects in *Lmna* mutant muscle cells are nucleus-intrinsic or arise from altered signaling pathways or other cytoplasmic changes in the mutant cells, we generated “hybrid” myofibers by combining wild-type and *Lmna* KO myoblasts prior to differentiation. Following myoblast fusion, these cells formed multinucleated myotubes and myofibers that contained both wild-type and *Lmna* KO nuclei with a shared cytoplasm (**Fig. 3e**). Importantly, in skeletal muscle cells, each nucleus is thought to provide mRNA transcripts for the nearby cytoplasm (referred to as myonuclear domain), so that the local RNA and protein content primarily stems from the nearest myonucleus^28^, and the genotype of each myonucleus can be determined by antibody staining against lamin A (**Fig. 3e**). We quantified the number of nuclei with chromatin protrusions and compared genetically identical nuclei (e.g., wild-type or *Lmna* KO) from hybrid and isogenic control myofibers after ten days of differentiation (**Fig. 3f**). Hybrid myofibers comprising ∼80% wild-type nuclei and ∼20% *Lmna* KO nuclei appeared healthy. Nonetheless, *Lmna* KO nuclei within the hybrid myofibers showed the same relative frequency of chromatin protrusions as nuclei from isogenic *Lmna* KO myofibers (**Fig. 3f**). Conversely, wild-type nuclei in hybrid fibers lacked chromatin protrusions and were thus not adversely affected by the presence of *Lmna* KO nuclei in the shared cytoplasm (**Fig. 3f**). These results indicate that the defects in nuclear structure are intrinsic to the *Lmna* mutant myonuclei and not due to impaired muscle fiber health, altered cytoplasmic signaling, or changes in the cytoplasmic architecture in *Lmna* mutant muscle fibers.

### *Lmna* mutant myofibers exhibit extensive NE rupture *in vitro*

Physical compression by cytoskeletal forces can result in NE rupture, with depletion of lamins exacerbating the frequency of NE rupture^13, 29–33^. To examine whether the reduced nuclear stability seen in *Lmna* mutant muscle cells (**Fig. 2**) leads to NE rupture in *Lmna* mutant myofibers, we modified primary myoblasts to co-express a fluorescent NE rupture reporter, consisting of a green fluorescent protein with a nuclear localization signal (NLS-GFP)^13^ and fluorescently labeled histone (H2B-tdTomato). NLS-GFP is normally localized to the nucleus, but rapidly spills into the cytoplasm upon loss of nuclear membrane integrity and is then gradually reimported into the nucleus after the nuclear membrane has been repaired^13^. *In vitro* differentiated *Lmna* KO myotubes frequently exhibited NE ruptures (**Fig. 4a**, Suppl. Movie 7), which were absent in wild-type controls. To investigate NE rupture in more detail, we stably modified primary myoblasts with another fluorescent NE rupture reporter, cGAS-mCherry, which accumulates at the rupture site^13^ (**Fig. 4b**). Unlike the transient cytoplasmic NLS-GFP signal, however, the cGAS-mCherry accumulation persists even after the NE has been repaired^13, 29^. Wild-type myotubes had no detectable accumulation of cGAS-mCherry (**Fig. 4c**). In contrast,

**Figure 4.**
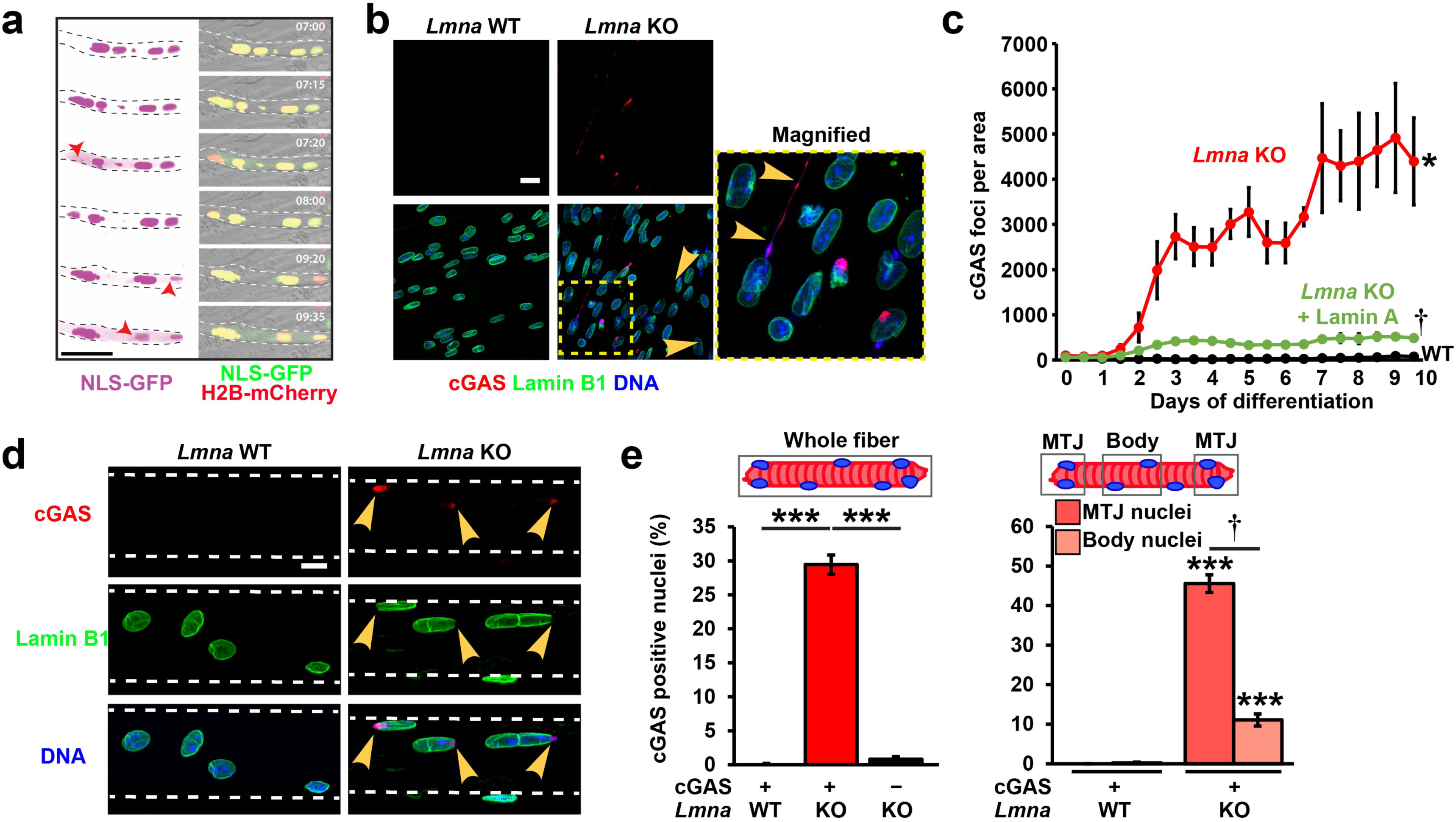
*Lmna* mutant myonuclei undergo NE rupture *in vitro* and *in vivo*. (**a**) Representative time-lapse image sequence of NE rupture in *Lmna* KO myonuclei. Red arrowheads mark two nuclei that undergo NE rupture, visibly by transient loss of NLS-GFP from the nucleus. Scale bar: 50 µm for all images. (**b**) Representative images of the accumulation of cGAS-mCherry at the sites of NE rupture in *Lmna* KO myonuclei at day 5 of differentiation. Scale bar: 20 µm. (**c**) Quantification of cGAS-mCherry foci formation per field of view during myofiber differentiation in *Lmna* WT, *Lmna* KO, and *Lmna* KO cells expressing ectopic lamin A, expressed. *n* = 3 independent experiments. *, *p* < 0.05 vs. *Lmna* WT. †, *p* < 0.01 vs. *Lmna* KO. (**d**) Representative maximum intensity projection images of single muscle fibers from *Lmna* WT and *Lmna* KO mice expressing a cGAS-tdTomato NE rupture reporter, showing accumulation of cGAS-tdTomato at the site of nuclear NE in *Lmna* KO muscle fibers. Scale bar: 10 µm. **(e**) Quantification of the percentage of myonuclei positive for cGAS-tdTomato foci in isolated muscle fibers from *Lmna* WT and *Lmna* KO mice expressing the cGAS-tdTomato transgene (cGAS+) or non-expressing littermates (cGAS–). The latter served as control for potential differences in autofluorescence. Analysis performed for whole fiber (left) and by classification of nuclei located at the MTJ or within the body of the fiber (right). *n* = 5-8 mice per genotype, with 5 fibers per animal. ***, *p* < 0.001 *vs*. *Lmna* WT. †, *p* < 0.01 *vs*. nuclei in the muscle body.

*Lmna* KO myotubes displayed a progressive increase in the number of nuclear cGAS-mCherry foci during differentiation, starting around day two, which could be rescued by ectopic expression of wild-type lamin A (**Fig. 4c**). *Lmna* N195K showed intermediate levels of NE rupture (Suppl. Fig. S7A), whereas *Lmna* H222P myotubes had cGAS-mCherry accumulation comparable to wild-type controls (Suppl. Fig. S7b), consistent with the milder defects in nuclear stability in the *Lmna* H222P mutant cells (**Fig. 2b,c**; Supp. Fig. S4a).

### Lamin A/C-deficient myofibers experience extensive NE rupture *in vivo*

To test whether NE rupture occurs within *Lmna* KO muscle *in vivo*, we generated transgenic mice that express a fluorescent cGAS-tdTomato nuclear rupture reporter and crossed these mice into the *Lmna* mutant mouse models. Single hindlimb muscle fibers isolated from *Lmna* KO offspring expressing the cGAS-tdTomato reporter revealed a large fraction of myonuclei with cGAS-tdTomato foci, which were absent in both wild-type littermates expressing the cGAS-tdTomato reporter and in *Lmna* KO mice that were negative for the cGAS-tdTomato reporter (**Fig. 4d-f**). Within *Lmna* KO muscle fibers, the frequency of NE rupture was significantly higher at the MTJ than in the myofiber body nuclei (**Fig. 4e**; Suppl. Fig. S8), consistent with the increased frequency of chromatin protrusions in the MTJ myonuclei. The amount of cGAS-tdTomato accumulation scaled with disease severity: *Lmna* KO fibers had the highest amount of cGAS-tdTomato foci, *Lmna* N195K fibers had an intermediate amount, and *Lmna* H222P fibers had no nuclear cGAS-tdTomato accumulation, closely matching the *in vitro* data (Suppl. Fig. S7c). As an independent approach to detect loss of nuclear-cytoplasmic compartmentalization in muscle fibers, we analyzed the intracellular localization of endogenous heat shock protein 90 (Hsp90), which is typically excluded from the nucleus in healthy muscle fibers^34^. In our *in vitro* assays, Hsp90 was cytoplasmic in wild-type myofibers, whereas all three *Lmna* mutant models had increased nuclear Hsp90 levels during myoblast differentiation (Suppl. Fig. S9a,b). Similarly, muscle fibers isolated from *Lmna* KO mice, but not wild-type littermates, had a significant increase in nuclear Hsp90 (Suppl. Fig. S9c,d), confirming the occurrence of NE rupture *in vivo*. Taken together, these findings indicate widespread NE rupture in severe cases of laminopathic skeletal muscle.

### *Lmna* KO myonuclei have increased levels of DNA damage *in vitro* and *in vivo*

Recent studies found that nuclear deformation and NE rupture can cause DNA damage in migrating cells^13, 29, 35^. To investigate whether chromatin protrusions and NE rupture can similarly lead to DNA damage in muscle cells, we quantified DNA damage in differentiating primary myoblasts by staining for γH2AX, a marker for double stranded DNA damage^36^. Both *Lmna* KO and wild-type myoblasts had elevated levels of DNA damage at the onset of differentiation (**Fig. 5a,b**), consistent with previous reports that show the transition from myoblasts to myotubes is associated with a transient increase in γH2AX levels^37, 38^. However, while γH2AX levels in wild-type myotubes subsequently decreased and then remained stable at low levels, the fraction of myonuclei with severe DNA damage in the *Lmna* KO cells continued to increase from five to ten days post-differentiation, with nearly 20% of *Lmna* KO myonuclei exhibiting severe DNA damage at day ten (**Fig. 5b**). Consistent with the increased DNA damage, *Lmna* KO myotubes exhibited significantly increased activity of the DNA-dependent protein kinase, DNA-PK (**Fig. 5c**), one of the major DNA damage response pathways in post-mitotic cells^36^. Single muscle fibers isolated from *Lmna* KO mice similarly contained many myonuclei with extensive γH2AX staining (**Fig. 5d,e**) and increased DNA-PK activity, especially at the MTJ, where over 10% of nuclei show very high intensity staining (**Fig. 5f,** Suppl. Fig S10), confirming the presence of extensive DNA damage in *Lmna* KO muscle fibers *in vivo*. The *Lmna* N195K single muscle fibers showed similar high levels of DNA damage compared to *Lmna* KO mice, whereas the *Lmna* H222P single muscle fibers showed an increase only in mid and low levels of DNA damage, consistent with the milder disease severity and other phenotypic markers (**Fig. 5e**). In contrast, muscle fibers isolated from wild-type mice contained only low levels of DNA damage (**Fig. 5d,e**). Of the myonuclei with extensive γH2AX staining, 82% also had chromatin protrusions, suggesting a link between physical damage to the nucleus and DNA damage (Suppl. Fig. S11).

**Figure 5.**
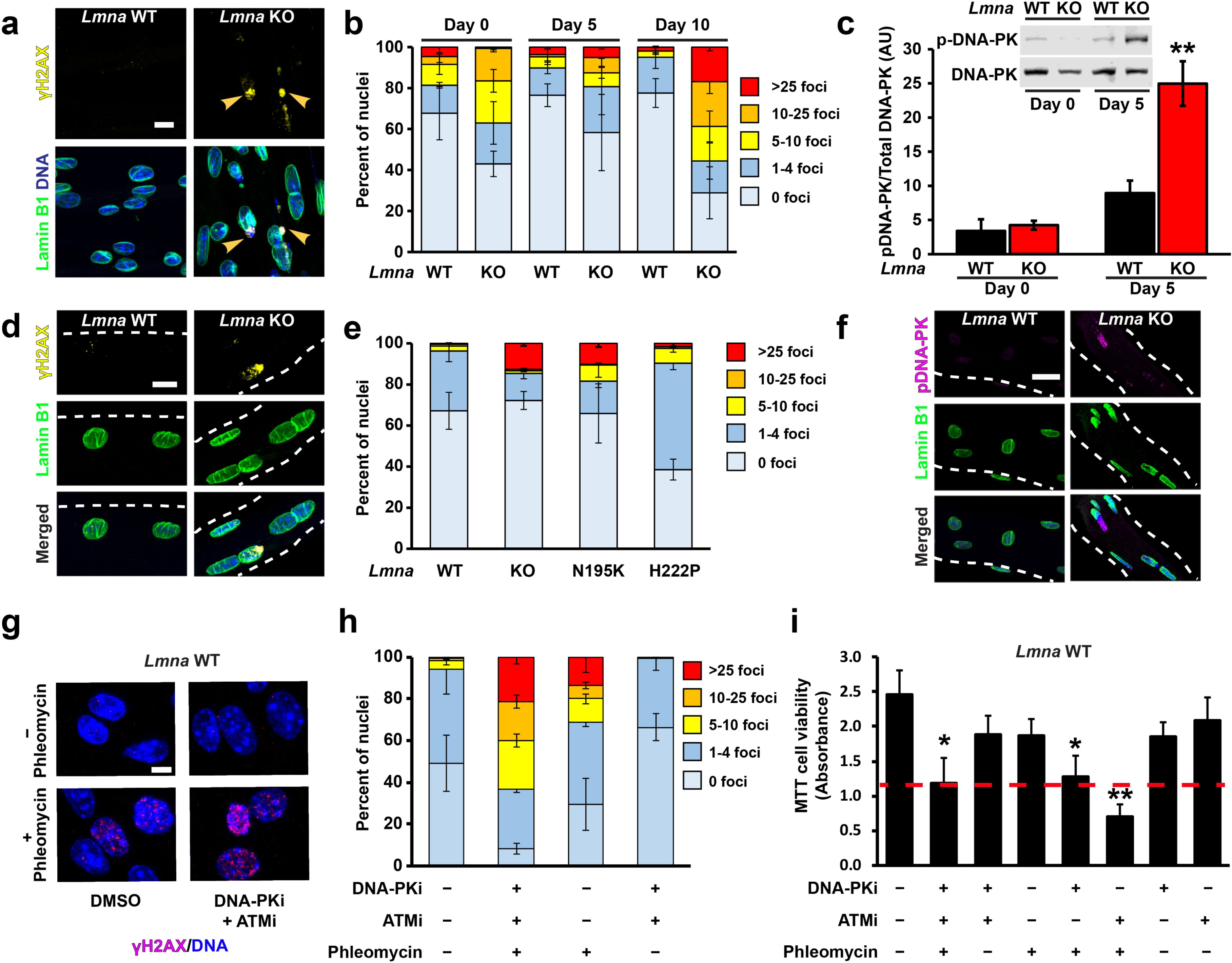
*Lmna* KO mice have increased DNA damage in myonuclei *in vitro* and *in vivo*. (**a**) Representative images of γH2AX foci, a marker of a double-stranded DNA break, in *Lmna* KO myonuclei. Arrowheads indicated γH2AX foci at the sites of chromatin protrusions. Scale bar: 10 µm. (**b**) Quantification of the extent of DNA damage based on the number of γH2AX foci per nucleus during myofiber differentiation. *Lmna* KO myonuclei show a progressive increase in the amount of severe DNA damage during myofiber differentiation. *n* = 3 independent cell lines per genotype. (**c**) Quantification of DNA-PK activity in *Lmna* WT and *Lmna* KO myotubes at day 5 of differentiation by probing for the phosphorylation of DNA-PK at S2053, an autophosphorylation specific site. *n* = 3 lysates from independent cell lines. **, *p* < 0.01 vs. *Lmna* WT. (**d**) Representative images of γH2AX foci in isolated single muscle fibers from *Lmna* WT and *Lmna* KO mice. Scale bar: 10 µm. (**e**) Quantification of the extent of DNA damage based on the number of γH2AX foci per nucleus in isolated single fibers. *n* = 3-5 mice per genotype in which 5 fibers are imaged per animal. (**f**) Representative image of p-DNA-PK (S2053) in isolated muscle fibers from *Lmna* WT and *Lmna* KO mice. Scale bar: 20 µm. (**g**) Representative image of γH2AX foci following treatment with phleomycin, with or without DNA-PK + ATM inhibition. Scale bar: 10 µm. (**h**) Quantification of the extent of DNA damage based on the number of γH2AX foci per nucleus for *Lmna* WT cells treated with phleomycin, with or without DNA-PK + ATM inhibition. Inhibiting DNA damage repair in the presence of phelomycin results in a significant increase in DNA damage above that of phelomycin alone. *n* = 3 independent experiments per condition. Quantification of cellular viability in *Lmna* WT myofibers using MTT assay following DNA damage induction with phleomycin, with and without concurrent treatment with DNA-PKi (NU7441) and/or ATMi (KU55933). *n* = 3 independent experiments per condition. **, *p* < 0.01 vs. untreated control; *, *p* < 0.05 vs. untreated control. Dashed red line indicates the corresponding quantity of the *Lmna* KO untreated control.

To determine whether the accumulation of DNA damage during *in vitro* differentiation of *Lmna* KO myoblasts was caused by progressive new nuclear damage or defects in DNA damage repair, which had been reported in several progeroid laminopathies^39–41^, we subjected *Lmna* KO and wild-type myofibers to a pulse of gamma irradiation and monitored γH2AX levels at 3, 6, and 24 hours post-treatment. Consistent with previous studies^42^, irradiation resulted in a rapid increase in the number of γH2AX foci at 3 hours that then gradually resolved and returned to baseline by 24 hours post irradiation (Suppl. Fig. S12). Notably, following irradiation, *Lmna* KO myofibers displayed a DNA damage profile nearly identical to wild-type controls, suggesting that their ability to repair DNA damage is not significantly impaired, and the accumulation of DNA damage in the myotubes is more likely due to new incidents of damage.

### Accumulation of DNA damage correlates with myofiber death

To test whether accumulation of DNA damage is sufficient to explain the progressive decline in myofiber viability in the *Lmna* KO cells, we subjected *Lmna* KO and wild-type myofibers to repeated treatments of phleomycin, a radiation mimetic agent, in conjuction with inhibition of DNA damage repair with NU7441, a DNA-PK-specific inhibitor, and/or KU55933, an ATM-specific inhibitor (**Fig. 5g**). Combined treatment with phleomycin and DNA damage repair inhibition (by one or both inhibitors in combination) resulted in an accumulation of DNA damage and loss of viability in wild-type myofibers comparable to that observed in untreated *Lmna* KO cells (**Fig. 5h,i;** Suppl. Fig.13). In contrast, phleomycin, alone or in combination with DNA damage repair inhibition, did not further reduce viability in *Lmna* KO myofibers (Suppl. Fig.13), suggesting that the preexisting DNA damage in these cells is already sufficient to drive myofiber decline. Since different types of DNA damage can elicit distinct repair responses in post-mitotic muscle cells^43^, the DNA damage and ensuing repair response associated with NE rupture may differ from that induced by either phleomycin or irradiation. Thus, it cannot be ruled out that the *Lmna* KO cells may exhibit some defects in DNA damage repair, or that additional mechanisms contribute to the pathogenesis in the *Lmna* mutant cells.

### Nuclear damage in *Lmna* KO myofibers can be prevented by microtubule stabilization

We surmised that NE ruptures in *Lmna* mutant myofibers resulted from cytoskeletal forces acting on mechanically weak myonuclei, and that reducing mechanical stress on the nuclei would decrease nuclear damage. In striated muscle cells, the microtubule network remodels significantly upon differentiation to form a cage-like structure around the myonuclei^44^ (Suppl. Fig. S14), in large part due to the redistribution of centrosomal proteins at the NE^45^. To test if stabilizing this microtubule network and thereby reinforcing myonuclei can reduce chromatin protrusions and NE rupture, we treated *in vitro* differentiated myoblasts with low doses of the microtubule stabilizing drug, paclitaxel. Here, we focused on the *Lmna* KO model, which showed the most severe nuclear defects. The microharpoon assay confirmed that microtubule stabilization reinforced *Lmna* KO nuclei in differentiated myofibers and significantly reduced nuclear deformation in response to cytoplasmic force application (**Fig. 6a,b**, Suppl. Movie 8). Furthermore, paclitaxel treatment significantly reduced the percentage of nuclei with chromatin protrusions (**Fig. 6c**) and the incidence of NE rupture detected with the cGAS-mCherry reporter in the *Lmna* KO cells (**Fig. 6d**), suggesting that nuclear damage indeed arises from mechanical stress on the myonuclei and can be prevented by mechanically stabilizing the nuclei.

**Figure 6.**
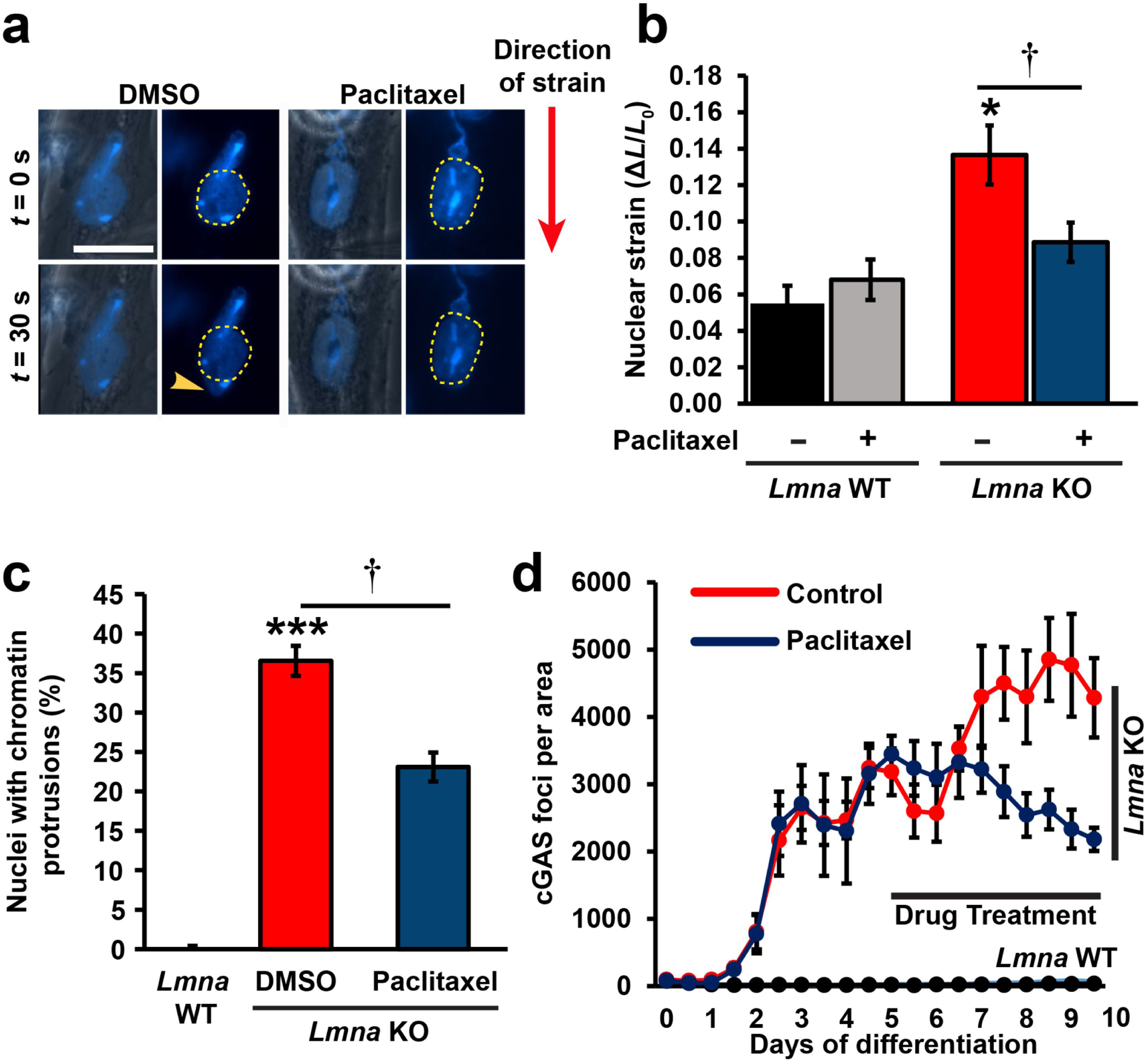
Mechanical reinforcement of *Lmna* KO myonuclei by microtubule stabilization reduces nuclear damage. (**a**) Representative image of nuclear deformation following microharpoon in *Lmna* KO myotubes at day five of differentiation. Myotubes were treated for 24 hours with either paclitaxel or DMSO control. Yellow dotted line denotes the perimeter of the nucleus prior to strain application. Scale bar: 20µm. (**b**) Quantification of nuclear strain in *Lmna* WT and *Lmna* KO myofibers using microharpoon assay following 24 hours of treatment with 50 nM paclitaxel or DMSO vehicle control. *, *p* < 0.05 *vs*. *Lmna* WT. †, *p* < 0.05 *vs*. vehicle control. (**c**) Quantification of chromatin protrusions at day 7 of differentiation following treatment with the paclitaxel (50 nM) or DMSO starting at day 4. *n* = 3 independent experiments. ***, *p* < 0.001 *vs*. *Lmna* WT. †, *p* < 0.01 *vs*. vehicle control. (**d**) Quantification of cGAS-mCherry foci formation during 10 myofiber differentiation following treatment with paclitaxel (10 nM) or DMSO control, starting at day 5 of differentiation. *n* = 3 independent experiments.

### Kinesin-mediated nuclear movements are responsible for nuclear damage in *Lmna* KO myonuclei

The findings that nuclear damage occurred only during myoblast differentiation, when the cytoskeleton significantly remodels, and that microtubule stabilization significantly reduced the amount of nuclear damage in *Lmna* KO myofibers, suggest that the nuclear defects result from cytoskeletal forces acting on the mechanically weaker *Lmna* mutant myonuclei. Physical stress may be imparted to myonuclei via (1) actomyosin-mediated contractile forces, and/or (2) forces due to nuclear movements at various stages of muscle development, including myoblast migration and fusion^46^, microtubule-driven spacing^44, 47–54^, shuttling to the periphery of myofibers^55–57^, and anchoring at the fiber periphery and neuromuscular junctions^58^. Recently, actomyosin contraction was shown to be a primary driver of NE rupture in chick cardiac tissue, and this NE rupture was rescued by treatment with blebbistatin^59^. To address whether NE damage in myotubes was caused by actomyosin contractility, we treated *Lmna* KO and wild-type myotubes with nifedipine, a muscle-specific calcium channel blocker. Nifedipine treatment effectively abrogated myotube contraction, but did not reduce the frequency of chromatin protrusions and NE ruptures in *Lmna* KO myonuclei (Suppl. Fig. S15), indicating that actomyosin contractility is not required to induce NE damage. The differences in our results could stem from differences in contraction strength between *in vitro* and *in vivo* systems as well as from the differences between cardiac and skeletal muscle cells. Therefore, we focused our attention to cytoskeletal forces exerted on the nucleus during nuclear migration in differentiating myotubes.

Based on the timing of the onset of NE rupture (**Fig. 4c**) and a progressive increase in the length of the chromatin tethers with differentiation (Suppl. Fig. S16), we examined the role of microtubule driven myonuclear spreading during myofiber maturation^60, 61^. Time-lapse sequences of *Lmna* KO myoblasts expressing the NLS-GFP and/or cGAS-mCherry reporters revealed that NE rupture frequently occurred when myonuclei were moved along the length of myotubes by microtubule-associated motors (**Fig. 7a,** Suppl. Movie 9). Thus, we reasoned that inhibiting nuclear movement should prevent nuclear damage in *Lmna* KO myofibers. Supporting this hypothesis, depletion of Kif5b (Suppl. Fig 17a), a subunit of kinesin-1 that is required for nuclear migration in myotubes ^60–62^, nearly abolished chromatin protrusions (Suppl. Fig. S17b), NE rupture (**Fig. 7b,c**), and high levels of DNA damage in the *Lmna* KO myotubes (Suppl. Fig. 17c,d). These findings indicate that nuclear movement by microtubule-mediated forces are sufficient to cause nuclear damage in *Lmna* mutant myofibers that correlates with increased DNA damage in these cells.

**Figure 7.**
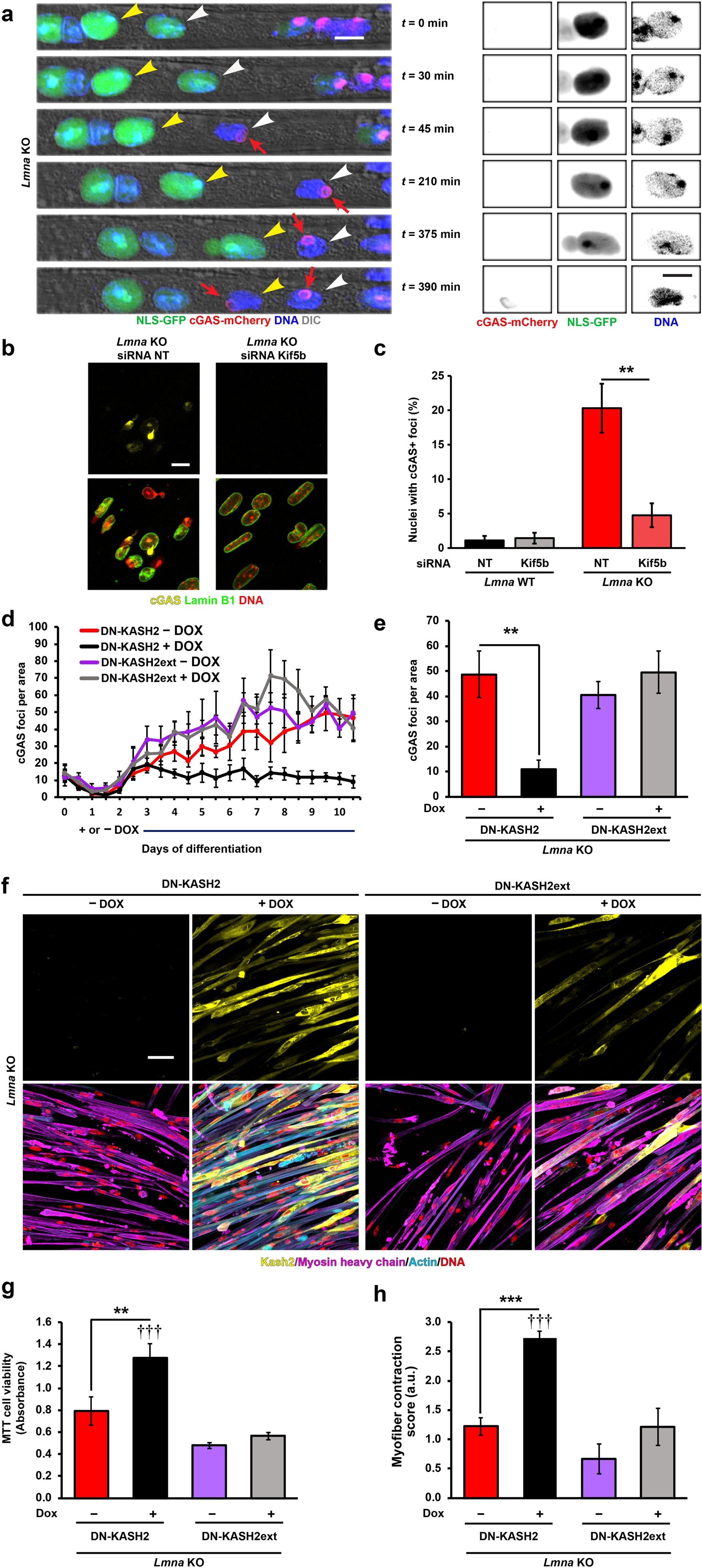
Reducing cytoskeletal forces on myonuclei prevents NE rupture and improves viability and contractility in *Lmna* KO myotubes. (**a**) Representative time-lapse image sequence of NE rupture in *Lmna* KO myonuclei during nuclear migration at day five of differentiation. White and yellow arrowheads mark two individual nuclei that undergo NE rupture, visible by transient loss of NLS-GFP from the nucleus and stable accumulation of cGAS-mCherry at the site of rupture (red arrow). Images on the right show close-ups of the nucleus marked with a yellow arrowhead. Scale bar: 10µm for all images. (**b**) Representative images of cGAS-mCherry accumulation in *Lmna* KO cells treated with either non-target control siRNA (siRNA NT) or siRNA against Kif5b. Scale bar: 20 µm (**c**) Quantification of the number of *Lmna* KO myonuclei positive for cGAS-mCherry foci following treatment with either non-target siRNA (siRNA NT) or siRNA against Kif5b. *n* = 3 independent experiments, in which a total of 911-1383 nuclei per condition were quantified. **, *p* < 0.01 vs. NT. (**d**) Quantification of the number of cGAS-mCherry foci in *Lmna* KO expressing either DN-KASH or DN-KASHext, treated with or without 1 µM doxycycline (DOX) starting at day 3 until day 10 of differentiation. *n* = 3 independent experiments per condition. (**e**) Quantification of the number of cGAS-mCherry foci at 10 days of differentiation as shown in (d). **, *p* < 0.01 vs. DN-KASH L DOX. *n* = 3 independent experiments per condition. (**f**) Representative images of *Lmna* KO expressing either DN-KASH or DN-KASHext, with or without 1 µM doxycycline (DOX), and immunofluorescently labeled for Myosin Heavy Chain, Actin and DAPI showing increased cell area and enhanced sarcomeric staining in the cells expressing DN-KASH + DOX. Scale bar: 50 µm (**g**) Quantification of cell viability following DN-KASH or DN-KASH-ext treatment in *Lmna* KO cells using the MTT assay. *n* = 6 per condition from 3 independent experiments. **, *p* < 0.01 vs. DN-KASH L DOX; †††, *p* < 0.001 vs DN-KASH L DOX and DN-KASH + DOX. (**h**) Quantification of myofiber contraction following DN-KASH or DN-KASHext treatment in *Lmna* KO cells using based on the percent of contractile fibers . *n* = 4 independent experiments per condition. ***, *p* < 0.001 vs. DN-KASH L DOX; †††, *p* < 0.001 vs DN-KASH L DOX and DN-KASH + DOX.

### Reducing the mechanical forces acting on *Lmna* KO myonuclei improve myofiber function and viability

If mechanically induced NE rupture and DNA damage are causative for the decline in viability and contractility in the *Lmna* KO myofibers, then reducing the cytoskeletal forces on the myonuclei should improve myofiber health. Since depletion of Kif5b has detrimental long-term effects on myofiber function^63^, we utilized a system where the Linker of Nucleoskeleton and Cytoskeleton (LINC) is disrupted through expression of a dominate negative GFP-KASH2 (DN-KASH) protein^64^ (Suppl. Fig. 17a). As a control, we generated a similar construct that contained a double alanine extension (DN-KASHext). The DN-KASHext construct still targets to the NE, but cannot disrupt the LINC complex^64^, as confirmed by the retention of nesprin-1 at the NE (Suppl. Fig. 18a). The DN-KASH constructs were expressed under the control of an inducible promotor that allowed for controlling the onset of LINC complex disruption following the initial fusion events of myogenesis (Suppl. Fig. S18b). This experimental approach reduced force transmission from the cytoskeleton to the nucleus and limited nuclear spreading in the DN-KASH cells, but not the DN-KASHext cells, similar to the depletion of Kif5b (Suppl. Fig. 18b). Expression of the DN-KASH constructs had no effect on myofiber contractility or viability in *Lmna* WT myofibers (Suppl. Fig. S18c,d; Suppl. Movies 10-13). Expression of the DN-KASH construct, but not the DN-KASHext, starting on day three of differentiation significantly reduced the number of chromatin protrusions and incidence of NE rupture in *Lmna* KO cells (**Fig. 7d,e,** Suppl. Fig. S18e). Importantly, this reduction in nuclear damage was accompanied by a reduction in the fraction of cells with severe DNA damage (Suppl. Fig. 18f) and substantially improved myofiber contractility and viability (**Fig. 7f-h;** Suppl. Movies 14-17). These findings further support our model that mechanical forces induce NE rupture and DNA damage in *Lmna* mutant myofibers, leading to myofiber dysfunction and death.

### Skeletal muscle biopsies from patients with *LMNA*-associated muscular dystrophy show DNA damage

Finally, to corroborate the findings from the *in vitro* and *in vivo* mouse models of striated muscle laminopathies in a clinically relevant context, we examined skeletal muscle biopsy samples from humans with *LMNA*-related muscular dystrophies and age-matched controls (**Table 1**). Muscle cryosections were immunofluorescently labeled for 53BP1, an established marker for DNA double strand breaks^65, 66^, and myofiber nuclei were identified based on labeling for DNA, actin, and dystrophin (**Fig. 8a,b**, Suppl. Fig. S19). Intriguingly, tissues from individuals with the most severe forms of the muscular dystrophy, i.e., those with early childhood and juvenile onsets, had significantly increased DNA damage compared to age-matched controls (**Fig. 8c-e**), thus closely mirroring the findings in the most severe laminopathy mouse models **(Fig. 5e).**

**Figure 8.**
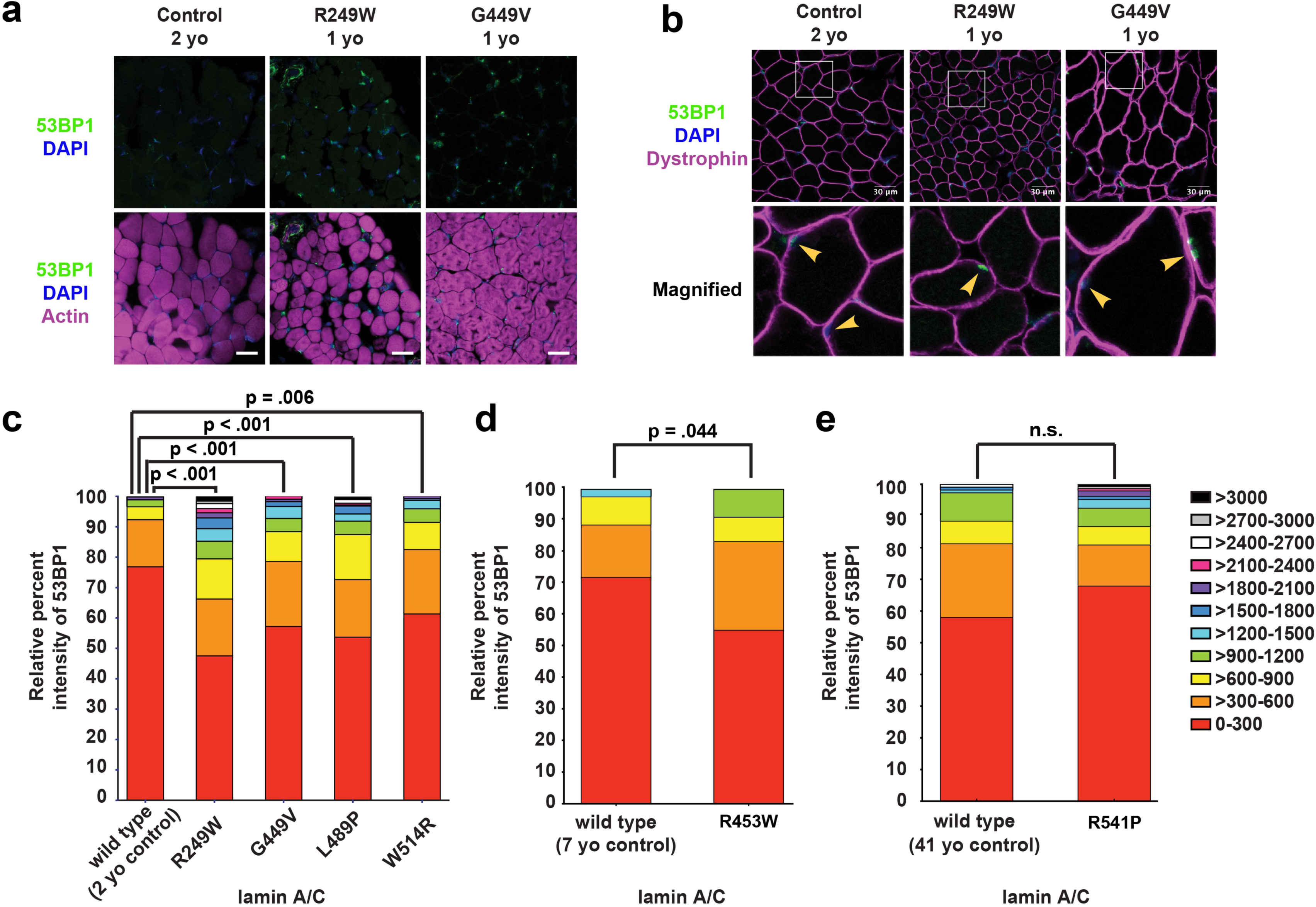
Human muscle biopsy tissue from individuals with *LMNA* muscular dystrophy showed increased 53BP1 staining. Cryopreserved human muscle biopsy tissue from individuals with *LMNA* muscular dystrophy and age-matched controls was sectioned and stained with either (**a**) anti-53BP1, DAPI, and phalloidin, or (**b**) anti-53BP1, DAPI, and dystrophin. Yellow arrowheads denote nuclei within muscle fibers, identified by anti-dystrophin immunolabeling of the muscle fiber membrane. Each muscular dystrophy patient possesses a *LMNA* mutation that results in a single amino acid substitution (Table 1). The *LMNA* mutations cause reduced fiber size, abnormally shaped fibers, and increased nuclear 53BP1 staining. Scale bar: 30 µm. (**c-e**) Patients were stratified based on the age at to time of muscle biopsy. The nuclear intensity values of 53BP1 were binned into 11 categories based on the level of intensity (color coding on the right). The X-axis represents the mutant lamin A/C expressed in individuals with an *LMNA* mutation and the age-matched control samples expressing wild-type lamin A/C. The Y-axis represents the relative percent intensity of 53BP1 staining quantified using ImageJ. Approximately 200 – 300 nuclei of each genotype were used for the quantification.

**Table 1.** Description of *LMNA* patients and muscle biopsy samples used for immunofluorescence analysis. n.a.; not available

| **Nucleotide change** | **Amino acid substitution** | **Age on onset** | **Age of diagnosis (years)** | **Age when muscle biopsy obtained (years)** | **Tentative diagnosis/comments** | **Reference** |
| --- | --- | --- | --- | --- | --- | --- |
| 745C>T | R249W | 3 months | <1 | 2 | Congenital MD, early onset with dropped head, unable to sit unassisted at age 3 years | unpublished |
| 1346G>T | G449V | 6 months | 3 | 1 | Congenital MD, non-ambulatory at age 13 years | 75, 105 |
| 1466T>C | L489P | <2 | 7 | 2 | Congenital MD , moderately severe at age 2 years | 75, 105 |
| 1540T>C | W514R | 2-3 yo | 5 | 3 | Mild to moderate myopathy at age 14 years | 75, 105 |
| 1357C>T | R453W | n.a. | 6 | 10 | Mild myopathy at age 5.5 years | unpublished |
| 1622G>C | R541P | n.a. | <52 | 52 | n.a. | unpublished |

## DISCUSSION

The mechanisms by which *LMNA* mutations result in skeletal muscle-specific diseases, such as AD-EDMD and *LMNA*-CMD, has long puzzled researchers and clinicians, presenting a major hurdle in the development of effective treatment approaches. Here, we present comprehensive new evidence in support of the ‘mechanical stress’ hypothesis. By using three mouse models of striated muscle laminopathies with varying onset and severity, we systematically assessed the frequency of nuclear defects, including chromatin protrusions, NE rupture, and DNA damage, in muscle fibers *in vitro* and *in vivo,* in a uniform genetic background. Our studies revealed a striking correlation between the nuclear defects observed during *in vitro* myoblast differentiation, myonuclear defects in isolated muscle fibers, and disease onset and severity in the mouse model. Notably, analysis of muscle biopsy tissue from individuals with *LMNA* muscular dystrophies showed a similar trend, with samples corresponding to the most severe forms of muscular dystrophy having the largest fraction of myonuclei with DNA damage. The paucity of available human muscle samples and effects of genetic background in these diseases, limit the conclusions that can be made.

Our findings of NE rupture are consistent with reports of NE damage and intrusion of cytoplasmic organelles into the nucleoplasm in skeletal muscle fibers of patients with EDMD^67–70^, cardiac myocytes in *LMNA*-dilated cardiomyopathy patients^71, 72^, lamin A/C-deficient mice^14, 73^, and muscle and tendons of lamin-deficient fruit flies^74, 75^. Unlike those previous reports, however, we now provide detailed information on the extent, timing, and cause of the NE damage, revealing a striking correlation of NE rupture and disease severity.

Although increased DNA damage and altered DNA damage repair have been reported previously in *LMNA* mutant cells, those cases were linked to progeroid diseases, including Hutchinson-Gilford progeria syndrome (HGPS) and atypical Werner syndrome (AWS)^39–41, 76, 77^. Our findings demonstrate for the first time that cytoskeletal forces acting on myonuclei result in nuclear damage and DNA damage in skeletal muscle fibers *in vitro* and *in vivo*. The precise mechanisms by which NE damage and NE rupture could cause DNA damage and cell death remains to be elucidated. The DNA damage could arise from exposure of genomic DNA to cytoplasmic nucleases following NE rupture, or nuclear exclusion and efflux of DNA repair factors, as previously discussed in the context of confined cell migration^78–80^. The correlation between DNA damage and NE defects in our studies (Suppl. Fig. S11) suggests that the DNA damage is linked to mechanically induced NE defects, which is further supported by the finding that depletion of kinesin-1 and disruption of the LINC complex abolished NE defects and the most severe DNA damage in *Lmna* KO cells (**Fig. 7b-e**, Suppl. Fig. 17b-d, S18e,f). Given the known association between lamin A/C and the DNA damage response protein 53BP1^81^, loss of lamin A/C could also impair DNA damage repair efficiency, although this effect may be limited to proliferating cells and the associated replication stress^82–84^. In our experiments, post-mitotic *Lmna* KO myofibers exposed to DNA-damaging irradiation exhibited similar DNA damage repair dynamics as wild-type cells, indicating that the observed increase in DNA damage in *Lmna* KO myofibers is not caused by defective DNA damage repair. Nonetheless, it is possible that the efficiency of DNA damage repair decreases over the course of the differentiation, as the highest levels of DNA damage were found in myofibers at late stages of maturation. Such a decrease in DNA repair efficiency is consistent with work showing that satellite cells repair radiation-induced DNA double strand breaks more efficiently than their differentiated counterparts^85^, and reduced DNA damage repair could amplify the progressive DNA damage in differentiating myoblasts. In wild-type myofibers, repeated exposure to DNA damaging agents, when combined with DNA damage inhibition, was sufficient to induce cell death to the same extent as observed in untreated *Lmna* KO cells (**Fig. 5g-i**), demonstrating that accumulating DNA damage is sufficient to induce cell death even in post-mitotic cells such as myofibers.

The role of DNA damage response signaling in post-mitotic muscle function is an area of increasing interest. DNA damage results in rapid activation of DNA damage response pathways, including DNA-PK and ATM, which results in stabilization of p53, one of the primary DNA damage response pathway that can induce cell cycle arrest, senescence, and apoptosis.^86, 87^ The consequences of increased DNA damage response signaling in post-mitotic cells remain poorly characterized, but recent findings point to an intriguing role of DNA damage response pathways in skeletal and cardiac muscle. Increased activity of DNA-PK, one of the major DNA damage sensing pathways in interphase cells^36^, was recently linked to the age-related decline of metabolic, mitochondrial, and physical fitness of skeletal muscle cells^34^. Furthermore, cardiac-specific expression of *Lmna* D300N results in increased DNA damage and activation of p53 in a laminopathy mouse model, and cardiac specific deletion of the *Trp53* gene encoding p53 significantly improved the cardiac defects, although only marginally improved overall survival^77^. Consistent with increased p53 signaling caused by accumulating DNA damage, we found evidence of caspase-3 activation and a progressive loss in viability in *Lmna* KO and N195K myofibers (**Fig. 1e,f**). While the mechanisms controlling apoptosis in a post-mitotic tissue such as skeletal muscle are not well understood and remain controversial^88^, this type of cell death has been observed in myofibers during other muscle wasting conditions^89^, in *Lmna* E82K mutant ^90^ and heterozygous *Lmna*^+/–^ mouse hearts^91^, and recently in a cardiac-specific *Lmna* D300N mouse model^77^.

Although *Lmna* mutant myoblasts have mechanically weaker nuclei than wild-type myoblasts, we found that NE damage only arose during myoblast differentiation, when cytoskeletal forces acting on the myonuclei increase. Surprisingly, nuclear damage during *in vitro* myofiber differentiation was associated with kinesin-1 mediated nuclear movements, independent of actomyosin contractions (**Fig. 7b,c,** Suppl. Fig. 15, Suppl. Fig. 17b-d). Kinesin-1 applies localized point forces at the NE of skeletal myonuclei, either directly or through microtubules anchored at the NE through the LINC complex, to ensure correct nuclear positioning^61^. These forces acting on the weakened NE are likely sufficient to induce NE rupture in the *Lmna* mutant myonuclei, based on recent studies on lamin A/C-depleted cells subjected to precisely controlled tensile forces^92^. Furthermore, mutations in genes encoding lamins A/C and other NE proteins linked to muscular dystrophies could disrupt perinuclear cytoskeletal organization, including that of desmins^73^ and the perinuclear microtubule network, which helps to resist cytoplasmic strain and physically protect myonuclei, thereby further promoting nuclear damage in myofibers. At the same time, actomyosin contractility is likely to contribute to NE damage in muscle fibers *in vivo*, which can generate substantially higher forces than *in vitro* differentiated myofibers, and to NE rupture and DNA damage in cardiac myocytes^59^.

While we cannot exclude the possibility that altered cytoplasmic signaling and gene regulation pathways contribute to the increased NE rupture and DNA damage in *Lmna* mutant muscle cells^77, 93^, our data suggest that the damage is mechanically induced and nucleus-intrinsic, as depletion of Kif5b and disruption of the LINC complex substantially reduced DNA damage in *Lmna* KO myofibers (**Figs. 3e,f, 4f,g,** Suppl. Fig S18e,f). Our findings support a model in which cytoskeletal forces cause chromatin protrusions and NE ruptures in mechanically weakened *Lmna* mutant muscle cell nuclei, triggering DNA damage, which then leads to myofiber dysfunction and death in striated muscle laminopathies (Supp. Fig. 20). Given the diverse roles of lamins in cellular function, additional mechanisms may further contribute to the pathogenesis of laminopathies, and their specific contribution may depend on the particular mutation and cellular context.

Our experimental data provides novel mechanistic insights into the cellular processes that contribute to the development of striated muscle laminopathies, thereby informing future research efforts to target. Microtubule stabilization is one approach to correct the perturbed force balance and should be further explored as a therapy for striated muscle laminopathies. Paclitaxel was recently reported to improve cardiac conduction defects in *Lmna* H222P mice by restoring proper connexin 43 localization^94^. Here, we highlight an additional mechanism by which microtubule stabilization may mitigate damaging forces in striated muscle laminopathies. In addition, our work indicates that DNA damage is increased in mouse models of laminopathies, as well as human patients, and that targeting cell signaling pathways activated by DNA damage, such as DNA-PK^34^ and p53^77^ may provide a mechanism to ameliorate muscle wasting. Furthermore, our studies found that hybrid myofibers containing ∼20% *Lmna* KO and ∼80% wild-type nuclei were indistinguishable from their isogenic wild-type controls, suggesting that delivery of wild-type lamin A/C to a subset of myonuclei by gene delivery or stem cell therapy may be sufficient to rescue myofiber function. Further studies will need to identify the critical number of wild-type nuclei required to rescue cellular health in laminopathic tissue for therapeutic applications.

Beyond striated muscle laminopathies, insights gained from this work are highly relevant to other biological systems in which nuclei are exposed to physical stress from the cytoskeleton, such as in confined migration of cancer cells^95^, or intracellular nuclear positioning of polarized epithelial, or neuronal cells^96^. For example, in neuronal cells lacking functional lamin A^97^, defects in B-type lamins^98^ or other NE proteins (e.g., nesprin-1) may make these cells more susceptible to kinesin-mediated nuclear damage leading to neurodevelopmental defects (e.g., cerebellar ataxia). Taken together, these findings highlight a novel mechanism by which weakened myonuclei experience microtubule-mediated NE damage, leading to DNA damage and muscle dysfunction, potentially explaining the phenotypes seen in striated muscle laminopathies and a spectrum of other diseases caused by NE defects.

## ACKNOWLEDGEMENTS

The authors thank Colin Stewart for providing the *Lmna* KO and *Lmna* N195K mouse models, Edgar Gomes for help with the *in vitro* myocyte differentiation protocol, Alexandra Corbin for *in vivo* and *in vitro* protrusion analysis and *in vivo* Hsp90 localization analysis, Rebecca Mount for optimization of single fiber isolation and subsequent analysis, Daniel Huang for quantification of the *Lmna* N195K skeletal muscle fiber cross-sectional areas, Ern Hwei Hannah Fong for quantification of contraction scores, Francoise Vermeylen, Stephen Parry and Lynn Johnson from the Cornell Statistical Consulting Unit, and Katherine Strednak and the Cornell Center for Animal Resources and Education (CARE) for help in maintaining the *Lmna* mutant mice. Clinical data from *LMNA* individuals was provided by Katherine D. Mathews, M.D. (Vice Chair for Clinical Investigation, Director, Muscular Dystrophy Clinic, Director, Iowa Neuromuscular Program, Professor of Pediatrics – Neurology and Professor of Neurology). Technical assistance with human muscle immunohistochemistry was provided by Nicholas M. Shaw and Margaret R. Ketterer (U. Iowa). This work was supported by awards from the National Institutes of Health [R01 HL082792 and U54 CA210184], the Department of Defense Breast Cancer Research Program [Breakthrough Award BC150580], the National Science Foundation [CAREER Award CBET-1254846 and MCB-1715606], the Muscular Dystrophy Association [Development Award MDA603238 to T.J.K.] as well as generous gifts from the Mills family, a Fleming Postdoctoral Fellowship to T.J.K, and National Science Foundation Graduate Research Fellowships to A.J.E. [2013160437] and G.R.F. [2014163403]. Financial support from the Muscular Dystrophy Association (477283 to L.L.W); Burroughs Welcome Fund Collaborative Research Travel Grant (#1017502); University of Iowa Wellstone Muscular Dystrophy Cooperative Research Center (U54, NS053672 to S.A.M). This work was performed in part at the Cornell NanoScale Facility, a member of the National Nanotechnology Infrastructure Network, which is supported by the National Science Foundation [Grant ECCS-0335765].

## AUTHOR CONTRIBUTIONS

A.E., T.K., G.F., P.I., L.W., and J.L contributed to the conception and design of the work; A.E., T.K., G.F., P.I., J.P., S.I., S.M. contributed to data acquisition and analysis; A.E., T.K., G.F., L.W., G.B., J.L. contributed to the interpretation of data; A.E., T.K., G.F., J.L contributed to drafting the manuscript. All authors contributed to editing the manuscript.

## METHODS

### Animals

*Lmna* KO (*Lmna*^–/–^)^14^, *Lmna* H222P (*Lmna*^H222P/H222P^)^16^, and *Lmna* N195K (*Lmna*^N195K/N195K^)^15^ have been described previously. *Lmna*^+/–^, *Lmna*^H222P/+^, and *Lmna*^N195K/+^ mice were backcrossed at least seven generations into a C57-BL/6 line. For each mouse model, heterozygous mice were crossed to obtain homozygous mutants, heterozygous mice, and wild-type littermates. *Lmna* mutant mice were provided with gel diet (Nutri-Gel Diet, BioServe) supplement to improve hydration and metabolism upon onset of phenotypic decline. *Dmd*^mdx^ mice have been described previously^99^; mice were obtained from the Jackson Laboratory in a C57BL background and hemi- or homozygous animals were bred to produce all hemi- and homozygous offspring. All mice were bred, maintained and euthanized according to IACUC approved protocols. Data from wild-type littermate controls for *Lmna* KO, *Lmna* N195K and *Lmna* H222P showed no difference in any of the experimental outcomes between the different wild-type littermates, so wild-type data was combined into a single group unless otherwise specified. For both *in vivo* and *in vitro* studies, cells and or tissues were isolated from a single mouse and counted as a single replicate. All data are based on at least two independently derived primary cell lines for each genotype.

### NE rupture reporter mouse (cGAS/MB21D1-tdTom transgenic mouse)

To detect NE ruptures *in vivo,* we generated a transgenic mouse expressing FLAG tagged human cGAS^E225A/D227A^ fused to a tdTomato fluorescent protein (cGAS-tdTomato) under the control of the commonly used constitutive active CMV promoter. The cGAS mutations are in the magnesium-binding domain, abolishing the enzymatic activity and downstream production of interferon, while still remaining the ability to bind to genomic DNA. The mammalian expression cassette including promoter and terminator (CMV-3xFLAG-cGAS^E225A/D227A^-tdTomato-SV40polyA) was released from the expression vector, removing the prokaryotic domains. The purified linear DNA was then injected into the pronucleus of fertilized embryos collected from super-ovulated C57BL/6 mice and transplanted into pseudo-pregnant recipients. The resulting transgenic mouse model was used to cross into the *Lmna* KO background to generate 3×FLAG-cGAS^E225A/D227A^-tdTomato positive *Lmna* KO mice within two generations.

### Myoblast isolation

Cells were harvested from *Lmna* KO, *Lmna* N195K, *Lmna* H222P, and wild-type littermates between 3-5 weeks for *Lmna* KO mice, 4-6 weeks for *Lmna* N195K, and 4-10 weeks for *Lmna* H222P mice using a protocol adapted from^14^. With the exception of the *Lmna* KO myoblasts, these time-points were prior to the onset of disease phenotypes. Myoblasts from wild-type littermates were harvested at the same time. Muscles of the lower hindlimb were isolated, cleaned of fat, nerve and excess fascia, and kept in HBSS on ice until all mice were harvested. The muscles were digested in 4 ml:1 g of tissue wet weight in a solution of 0.5% Collagenase II (Worthington Biochemicals), 1.2 U/ml Dispase (Worthington Biochemicals), 1.25 mM CaCl_2_ (Sigma) in HBSS/25 mM HEPES buffer. Digestion was carried out in a 37°C water bath for a total time of 60 minutes. At 20 minutes intervals, digestion cocktails were removed and triturated 40 times with a 5 ml pipet. In the case of difficult to digest tissues, an extra 25% of 1% Collagenase II was added to the digestion after 40 minutes.

When tissues were fully digested, the reaction was quenched using equal volumes of DMEM supplemented with 10% fetal bovine serum (FBS) and 1% P/S (D10 media, Gibco). The cell suspension was strained through 70 and 40 μm filters (Greiner Bioscience) sequentially to remove undigested myotube fragments and tendon. The cell suspension was centrifuged at 800 × g for 5 minutes and washed with 8 ml of D10 media for a total of four times. Cells were then resuspended in primary myoblast growth media (PMGM; Hams F-10 (Gibco) supplemented with 20% horse serum and 1% penicillin/streptomycin and 1 µl/ml basic fibroblast growth factor (GoldBio)) and plated onto a 2% gelatin coated T25 flask. Cells were allowed to sit undisturbed for 72 hours. Capitalizing on the fact that myoblasts adhere much more weakly than fibroblasts, cells were passaged using PBS (calcium- and magnesium-free) instead of trypsin to purify the myoblasts. Cells were washed for 2-3 minutes at room temperature using a volume of PBS sufficient to coat the bottom of the flask and dislodged using manual agitation. When necessary, a 0.000625% trypsin solution was used to aid in the myoblast removal. Myoblasts were re-suspended in PMGM and re-plated onto gelatin coated flasks. This process was continued 3-4 times until pure myoblast cultures were achieved^100^. Cells were maintained in culture on gelatin coated flasks with media changes every other day. All experiments were carried out prior to passage 12. Each independent experiment was done on a different set of lamin mutant and wild-type littermates such that each independent experiment was sourced from a different animal to account for heterogeneity in phenotype.

### Myoblast differentiation

Myoblasts were differentiated according to a protocol modified from^17^. Coverslips for differentiation were prepared by first coating with CellTak (Corning) according to the manufacturer’s protocol and then coating with growth factor reduced Matrigel (Corning) diluted 1:100 with IMDM with Glutamax (Gibco). Pre-cooled pipette tips were used to avoid premature polymerization. Matrigel was allowed to polymerize at 37°C for 1 hour and the excess solution was aspirated. Primary myoblasts were seeded at a density of 55,000 cells/cm^2^ in PMGM. Cells were allowed to attach for 24 hours before being switched to primary myoblast differentiation media (PMDM) composed of IMDM with Glutamax and 2% horse serum without antibiotics. This timepoint was considered day 0. One day after the onset of differentiation, a top coat of 1:3 Matrigel:IMDM was added to the cells and allowed to incubate for 1 hour at 37°C. PMDM supplemented with 100 ng/ml agrin (R&D Systems) was added to the cells and henceforth replaced every second day. Cells were allowed to differentiate for a total of 0, 5, or 10 days.

### Plasmids and generation of fluorescently labeled cell lines

Each of the mutant myoblast lines were stably modified with lentiviral vectors to express the nuclear rupture reporter NLS-GFP (pCDH-CMV-NLS-copGFP-EF1-blastiS) and cGAS-mCherry (pCDH-CMV-cGAS^E225A/D227A^-mCherry2-EF1-blastiS). cGAS is a cytosolic DNA binding protein; we used a cGAS mutant (E225A/D227A) with abolished enzyme activity and interferon production, but that still binds DNA^101^ and serve as a NE rupture reporter^13^. For rescue experiments, *Lmna* KO cells were modified with human lamin A (pCDH-CMV-preLamin A-IRES-GFP-puro). To generate the DN-KASH and DN-KASHext constructs, GFP-KASH2 and GFP-KASH2ext were subcloned from previously published plasmids^64^ and inserted into an all-in-one doxycycline inducible backbone (pPB-tetO-GFP-KASH2-EIF1α-rtTA-IRES-Neo)^102^.

### Viral and Piggybac modification

Pseudoviral particles were produced as described previously^13^. In brief, 293-TN cells (System Biosciences, SBI) were co-transfected with the lentiviral-containing, packaging and envelope plasmids using PureFection (SBI), following manufactures protocol. Lentivirus-containing supernatants were collected at 48 hours and 72 hours after transfection, and filtered through a 0.45 µm filter. Cells to be transduced were seeded into 6-well plates so that they reached 50-60% confluency on the day of infection and transduced at most 2 consecutive days with the viral supernatant using the TransDux Max system (SBI). The viral solution was replaced with fresh culture medium, and cells were cultured for 72 hours before selection with 1 μg/mL of puromycin or 2 μg/mL blasticidin S for 2-5 days. After selection, cells were subcultured and maintained in their recommended medium without the continued use of selection agents. For PiggyBac modifications, myoblasts were transfected with 1.75 μg of the PiggyBac plasmid and 0.75 μg of a Hyperactive Transposase using the Lipofectamine 3000 reagent according to the manufacture’s guidelines.

### Extended imaging using incubator microscope

Long term imaging was performed using an Incucyte imaging system, which allows for incubator imaging to minimize the effects of humidity and CO_2_ changes. The differentiating cells expressing combinations of NLS-GFP and cGAS-mCherry were imaged using the Incucyte dual color filter module from day 0 to day 10, every 30-60 minutes with a 20× objective. Resulting images were analyzed using the Incucyte software, which performs fluorescence background subtraction using a top hat method and then subsequent thresholding. cGAS-mCherry cells were thresholded and then analyzed for increase in fluorescent foci over time to track the rate of increase in NE rupture or damage. NLS-GFP cells were used to investigate the frequency and presence of NE rupture. To verify the results obtained from the Incucyte, cells were fixed and stained with appropriate antibodies to evaluate DNA damage and NE rupture.

### Isolation of single muscle fibers

Single muscle fibers were harvested in a protocol adapted from Vogler et al.^103^. As previously described, fibers were isolated from the extensor digitorum longus (EDL) of male and female *Lmna* KO and wild-type litter mates at 5-6 weeks of age and *Lmna* H222P and wild-type litter mates were harvested at 6-8 weeks of age at 23-25 weeks of age in order to compare pre- and post-phenotype onset tissue^14, 16, 104^. Briefly, the EDL (extensor digitorus longus) and plantaris were isolated from the mouse and placed directly into a 1 ml solution of F10 media with 4,000 U/ml of Collagenase I (Worthington Biochemicals). The tissue was digested for 15-40 minutes depending on muscle size in a 37°C water bath with agitation by inversion every 10 minutes. The reaction was quenched by transferring the digestion mixture to 4 ml of PMGM. Single fibers were hand-picked from the digested tissue using fire polished glass Pasteur pipettes. When necessary, the tissue was further dissociated by manual pipetting with a large-bore glass pipet. Fibers were washed once in fresh media prior to fixation with 4% paraformaldehyde (PFA) for 15 minutes at room temperature and subsequent IF staining.

### Pharmacological treatments

For preliminary experiments, myoblasts were differentiated using the standard protocol and treated with pharmacological treatments starting at day 5 of differentiation. For chromatin protrusion studies, paclitaxel was administered to differentiated myotubes in two 24 hours bursts at day 4 and day 6-post differentiation with a 24 hour recovery in between. Myotubes were then fixed in 4% PFA at day 7 and stained with anti-lamin B and DAPI in order to quantify the percentage of myonuclei with chromatin protrusions. For long term studies using the cGAS reporter, myotubes were treated with 10 nM of paclitaxel starting at day 5 and then media was refreshed every day. To inhibit myotube contraction, cells were treated with 5 µM nifedipine starting at day 5 and then media was refreshed every day. For DNA damage induction and inhibitor experiments, cells were treated with 20µg/ml of phleomycin for a two-hour pulse on Day 3,4, and 5 of differentiation. Concurrently, cells were treated with NU7441 (2 μM), KU55933 (2 μM) starting at day 2 of differentiation through day 10 of differentiation. For induction of DN-KASH and DN-KASHext, cells were treated with 1 μM doxycycline.

### Biophysical assays

To evaluate nuclear deformability in high throughput, we designed and fabricated a microfluidic, micropipette aspiration device. The mask and wafers were produced in the Cornell NanoScale Science and Technology Facility (CNF) using standard lithography techniques. PDMS molds of the devices were cast using Sylgard 184 (Dow Corning) and mounted on coverslips using a plasma cleaner as described previously^13^. Three port entrances were made using a 1.2 mm biopsy punch. Pressures at the inlet and outlet ports were set to 1.0 and 0.2 psi (relative to atmospheric pressure, *P*_atm_), respectively, using compressed air regulated by a MCFS-EZ pressure controller (Fluigent) to drive single cells through the device. Myoblasts (∼5×10^6^ cells/mL suspended in 2 % bovine serum albumin (BSA), 0.2 % FBS and 10 μg/mL Hoechst 33342 DNA stain in PBS) were captured within an array of 18 pockets, and then forced to deform into 3 µm wide × 5 µm tall micropipettes. The selected pressures resulted in detectable nuclear deformations without causing significant damage to the cells (tested using propidium iodide staining). The remaining port was set to *P*_atm_ and outfitted with a handheld pipette to flush cells from the pockets at the start of each image acquisition sequence. Brightfield and fluorescence images were acquired every 5 seconds for a minimum of 60 seconds using an inverted microscope and 20×/NA 0.8 air objective. Nuclear protrusion length was calculated using a custom-written MATLAB program, made available upon request.

For the microharpoon studies, myoblasts were seeded in 35 mm glass bottom dishes and differentiated as previously described, except without the addition of a Matrigel top coat to allow microharpoon access. A Sutter P-97 micropipette puller was used to create microharpoons from borosilicate glass rods (Sutter; OD: 1.0 mm, ID: 0.78, 10 cm length) with tip diameters of ≈1 μm. Day 4 myotubes (*Lmna* KO and wild-type) were treated for 24 hours with either 50 nM Paclitaxel or the corresponding 0.1% DMSO. The following day, the microharpoon assay was performed as previously described by our laboratory^22^, with slight modifications to the pull parameters to accommodate myotubes. The microharpoon was inserted ≈5-7 μm from the edge of the nucleus and pulled 15 μm at a rate of 1 μm/s. Pull direction was always orthogonal to the long axis of the myofiber. Images were acquired at 40× (+1.6×) every 5 seconds. Nuclear strain and centroid displacement were calculated using a custom-written MATLAB program, made available upon request.

### siRNA treatment

siRNAs used were as follows: Kif5b#3 (target sequence 5′-CAGCAAGAAGTAGACCGGATA-3′; Qiagen SI00176050), Kif5b#4 (target sequence 5′-CACGAGCTCACGGTTATGCAA-3′; Qiagen SI00176057), and non-target (NT) negative control (ON-TARGETplus non-targeting pool, Dharmachon, D-001810-10). Myoblasts were seeded at a density of ∼15,000 cells per well in a 96-well glass bottomed dish containing a matrigel coating. Once adhered, the myoblasts were transfected twice, 48 hours apart, with siRNA for NT or Kif5b using Lipofectamine RNAiMAX at a concentration of 150 nM in PMGM. After 12 hours, the myoblasts were switched to PMDM and differentiated for 5 days.

### Immunofluorescence staining of mouse cells and tissues

Cells were fixed in pre-warmed 4% PFA at the appropriate time point(s) and washed with PBS. Cells were blocked and permeabilized with a solution of 3% BSA, 0.1% Triton-X 100 and 0.1% Tween (Sigma) for 1 hour at room temperature. Cells were stained with primary antibodies diluted in blocking solution according to Table 2 at 4°C overnight. Samples were washed with PBS and incubated for 1 hour at room temperature with 1:250 dilution of AlexaFluor antibodies (Invitrogen) and 1:1000 DAPI (Sigma). Single muscle fibers were stained using the same procedure in Eppendorf tube baskets with an increase in blocking solution Triton-X concentration to 0.25%.

### Human patient biopsy staining

Following diagnostic testing, muscle biopsies were stored at - 80° C and subsequently utilized for research following protocols approved by the corresponding IRB, with informed consent from all participants. Cryopreserved human quadriceps muscle biopsy tissue from *LMNA* muscular dystrophy individuals and age-matched controls were used for immunostaining as described ^105^. An anti-rabbit polyclonal 53BP1 antibody (Novus) and anti-dystrophin mouse monocolonal antibody (Mab7A10, U of Iowa Hospitals and Clinics Pathology Core) were used at 1:1000 and 1:20 dilutions, respectively. Texas Red labeled phalloidin (Invitrogen) was used at 1:400 dilution. Secondary antibodies were a goat anti-rabbit Ig Alexa 488 conjugate (Invitrogen) and a goat anti-mouse IgG rhodamine Red-X conjugated (Molecular Probes), both used at 1:500 dilution. Slides were imaged on a Zeiss 710 confocal microscope (University of Iowa Central Microscopy Facility). The intensity of nuclear anti-53BP1 staining was quantified using ImageJ. Analysis of the human muscle samples was performed in a double-blinded manner. A pathologist generated the 10 µm thick cryo-sections and coded the samples, which were stained, imaged, and quantified prior to decoding by an independent individual.

### Western analysis

Cells were lysed in RIPA buffer containing protease (cOmplete EDTA-Free, Roche) and phosphatase (PhosSTOP, Roche) inhibitors. Protein was quantified using Bio-Rad Protein Assay Dye and 25-30 µg of protein lysate was separated using a 4-12% Bis-Tris polyacrylamide gel using standard a standard SDS-Page protocol. Protein was transferred to a polyvinylidene fluoride (PVDF) membrane overnight at 4°C at a current of 40 mA. Membranes were blocked using 3% BSA in tris-buffered saline containing 0.1% Tween-20 and primary antibodies (Table 2) were diluted in the same blocking solution and incubated overnight at 4°C. Protein bands were detected using either IRDye 680LT or IRDye 800CW (LI-COR) secondary antibodies, imaged on an Odyssey® CLx imaging system (LI-COR) and analyzed in Image Studio Lite (LI-COR)

### Imaging acquisition

Cells on coverslips and mounted single muscle fibers were imaged with an inverted Zeiss LSM700 confocal microscope. Z-stack were collected using 20× air (NA = 0.8), 40× water-immersion (NA = 1.2) and 63× oil-immersion (NA = 1.4) objectives. Airy units for all images were set between 1 and 1.5. Epi-fluorescence images were collected on a motorized inverted Zeiss Observer Z1 microscope equipped with CCD cameras (Photometrics CoolSNAP EZ or Photometrics CoolSNAP KINO) or a sCMOS camera (Hamamatsu Flash 4.0). H&E histology images were collected on an inverted Zeiss Observer Z1 microscope equipped with a color CCD camera (Edmund Optics, EO-0312C).

### Image analysis

Image sequences were analyzed using ZEN (Zeiss), ImageJ, or MATLAB (Mathworks) using only linear adjustments uniformly applied to the entire image region. Region of interest intensities were extracted using ZEN or ImageJ. To quantify cleaved caspase-3 (i.e. active) area and myofiber health, maximum intensity protections were generated, which were then blinded to the observer. Cleaved caspase-3 area was calculated by thresholding of the caspase-3 and myosin heavy chain fluorescent signal and expressing the cleaved caspase-3 signal relative to the myosin heavy chain signal. Myofiber contractions were scored based on a minimum of 6 random fields of view per replicate using a blinded analysis according to the scales provided in Suppl. Fig. S3. To count the number of DNA protrusions, and DNA damage foci, confocal image stacks were three-dimensionally reconstructed and displayed as maximum intensity projections. Protrusions lengths were both counted and measured by the presence of DAPI signal beyond the lamin B rim of the nucleus. Aspect ratio was quantified based on a thresholded lamin B rim to avoid the confounding factor of the DNA protrusions outside the body of the nucleus. Nuclear rupture was detected by an increase of the cytoplasmic NLS-GFP signal, or the localization of cGAS-mCherry to the nucleus. For better visualization of NLS-GFP cells many of the fluorescent single color image sequences were inverted. Graphs were generated in Excel (Microsoft), and figures were assembled in Illustrator (Adobe). DNA damage was determined by counting H2AX foci and then binned based on foci number. If damage was so severe that individual foci could not be counted, these nuclei were placed in the >25 foci category. For Hsp90 quantification, average nuclear Hsp90 fluorescence intensity was determined from a single mid-nucleus z-plane image and normalized to the cytoplasmic intensity at two points immediately adjacent to the nucleus.

### MTT assay

Myoblasts, seeded in a 96-well plate and differentiated as previously described for 0, 5, or 10 days, were assayed for cell viability according to the manufacturer’s instructions (Promega, CellTiter 96 Non-Radioactive Cell Proliferation Assay). Fresh differentiation media was added two hours prior to the addition of 15 μL MTT 3-(4,5-dimethylthiazol-2-yl)-2,5-diphenyltetrazolium bromide dye. After incubation for 3 hours in MTT dye, 100 uL of Stop Solution was added to solubilize the formazan product (appears purple). Following overnight incubation at 37°C and 5% CO_2_, the absorbance of each well (measured at 590 nm) was analyzed using a microplate reader.

### Gamma irradiation

A pulse of gamma-irradiation (5 Gy) was administered to myotubes differentiating (5 days) in a 96 well plate. Non-irradiated controls, along with treated cells following 3, 6, or 24 hours of recovery, were PFA-fixed and stained with anti-γH2AX, anti-lamin B and DAPI. A custom macro was used to quantify the mean integrated density of nuclear γH2AX signal from maximum intensity projections of confocal z-stacks.

### Statistical analysis

Unless otherwise noted, all experimental results were taken from at least three independent experiments and *in vivo* data were taken from at least three animals. For data with normal distribution, we used either student’s t-tests (comparing two groups) or one-way ANOVA (for experiments with more than two groups) with post-hoc tests. When multiple comparisons were made, we adjusted the significance level using Bonferroni corrections. All tests were performed using GraphPad Prism. Micropipette aspiration data were natural log-transformed (Suppl. Fig. S4a) and analyzed by linear regression of the log-log data. In addition, data was analyzed with a multilevel model, in which the log-transformed protrusion length was the dependent variable in the model and the log-transformed time, genotype, and their interaction were treated as independent fixed effects. Variance from individual experiments and other effects were considered in the model as random effects. Post-hoc multiple comparisons test with Dunnett correction were performed to determine differences between *Lmna* mutant cells (*Lmna* KO, *Lmna* N195K, and *Lmna* H222P) and control cells (pooled wild-type). Analyses were carried out using JMP software. For human tissue samples, nuclei were binned into eleven categories based on their intensity of 53BP1 staining as measured with ImageJ (arbitrary units). The relative percent intensities of each bin were plotted as a histogram for each genotype. Since the intensity values were not in a normal distribution, the Kruskal-Wallis one-way analysis of variance (ANOVA), followed by the Dunn post hoc test, were used to determine if there was a significant difference in staining among the genotypes. The Mann-Whitney non-parametric test was used for comparisons that involved one sample and one age-matched control. Unless otherwise noted, * denotes *p* ≤ 0.05, ** denotes *p* ≤ 0.01, and *** denotes *p* ≤ 0.001. Unless otherwise indicated, error bars represent the standard error of the mean (SEM). The data that support the findings of this study are available from the corresponding author upon reasonable request.

### Data and code availability

The data that support the findings of this study are available from the corresponding authors upon reasonable request. MATLAB codes used for the microharpoon assay and micropipette aspiration analysis is available upon request.

