## Supplementary Materials & Figure Legends for "Mutant lamins cause nuclear envelope rupture and DNA damage in skeletal muscle cells"

**Supplemental Figures**

**Supplemental Figure 1**. ***Lmna* display varying degrees of muscular dystrophy**. (**a**) Quantification of the average myofiber cross-sectional area of *Lmna* WT and *Lmna* KO mice. (**b**) Relative frequency of myofiber cross-sectional area in *Lmna* WT and *Lmna* KO mice. ***, *p* < 0.001 vs. *Lmna* WT; *n* = 8-10 animals per genotype. (**c**) Quantification of the average myofiber cross-sectional area of *Lmna* WT and *Lmna* N195K mice. (**d**) Relative frequency of myofiber cross-sectional area in *Lmna* WT and *Lmna* N195K mice. **, *p* < 0.01 vs. *Lmna* WT; *n* = 11-12 animals per genotype. (**e**) Quantification of the average myofiber cross-sectional area of *Lmna* WT and *Lmna* H222P mice. (**f**) Relative frequency of myofiber cross-sectional area in *Lmna* WT and *Lmna* H222P mice. *n* = 3-4 animals per genotype.

**Supplemental Figure 2. *In vitro* differentiation results in mature myofibers**. Representative image of a striated myofiber containing a peripheral nucleus at day 10 of differentiation. Scale bar: 5 µm.

**Supplemental Figure 3.** ***Lmna* KO and *Lmna* N195K have reduced contractility and experience nuclear loss**. (**a**) Quantification of myofiber contraction at day 10 of differentiation. Fibers were assigned contraction scores from 0 (worst) to 4 (best) based on the percentage of cells that were visually contracting. **, *p* < 0.01 vs. *Lmna* WT, *, *p* < 0.05 vs. *Lmna* WT; *n* = 3-6 independent cell lines for each genotype. (**b**) Quantification of the change in nuclear number between day 5 and day 10 of differentiation. ***, *p* < 0.001 vs. *Lmna* WT *, *p* < 0.05 vs. *Lmna* WT; *n* = 3-6 independent experiments from 3 independent cell lines per genotype.

**Supplemental Figure 4**. **Micropipette aspiration analysis of *Lmna* mutant myoblasts and *Lmna* KO myoblasts ectopically expressing lamin A**. (**a**) Natural log transformation and plot of the micropipette aspiration data shown in Fig. 2b. The log-log data fits a linear regression model, in which all three *Lmna* mutants were significantly different (*p* < 0.001) from the wild-type controls. The slopes of the log-log data were not significantly different between the samples. A multilevel model including day-to-day variability confirmed that all three *Lmna* mutants were significantly different from the wild-type controls (*p* < 0.0001 for *Lmna* KO and *Lmna* N195K; *p* < 0.001 for *Lmna* H222P), although the statistical significance for the *Lmna* H222P myoblasts was lost when including additional variance components. (**b**) Representative immunofluorescence images of lamin A expression in *Lmna* WT, *Lmna* KO and *Lmna* KO cells ectopically expressing lamin A (*Lmna* KO + lamin A). (**c**) Measurement for nuclear deformation at 5 second intervals for *Lmna* WT, *Lmna* KO, and *Lmna* KO + Lamin A myoblasts during 60 seconds of aspiration. (**d**) Quantification of the nuclear deformation after 60 seconds of aspiration, showing that ectopic expression of lamin A significantly improves nuclear stiffness in *Lmna* KO myoblasts. *n* = 41-67 nuclei per genotype from 3 independent experiments. *n* = 62-73 nuclei per genotype from 3 independent experiments. ***, *p* < 0.001 *vs* *Lmna* WT cells. †, *p* < 0.01 *vs*. *Lmna* KO cells.

**Supplemental Figure 5.** ***Mdx* myoblasts have normal nuclear stiffness**. Measurement for nuclear deformation at 5 second intervals for *Lmna* WT and *mdx* myoblasts during 60 seconds of aspiration. *n* = 35-67 cells per condition.

**Supplemental Figure 6.** **Chromatin protrusions are surrounded by nuclear membranes containing emerin, with disturbed localization of nesprin-1 and nuclear pores**. (**a**) Representative immunofluorescence images for nesprin-1 and emerin in *Lmna* WT, *Lmna* KO, *Lmna* N195K and *Lmna* H222P myofibers at day 5 of differentiation. Blue and yellow arrows denote chromatin protrusions that are enriched with nesprin-1 and emerin, respectively. Scale bar: 20 µm. (**b**) Representative image of immunofluorescence detection of nuclear pore complexes (NPC) in *Lmna* KO myofibers at day 10 of differentiation. Scale bar: 10 µm.

**Supplemental Figure 7. Nuclear envelope rupture is increased in *Lmna* N195K myofibers *in vitro* and *in vivo***. Quantification of cGAS-mCherry nuclear envelope rupture reporter foci formation during 10 myofiber differentiation in *Lmna* N195K (**a**), *Lmna* H222P (**b**), and *Lmna* WT cells. (**c**) Quantification of the percentage of myonuclei positive for cGAS-tdTomato foci in isolated muscle fibers from *Lmna* WT, *Lmna* KO, *Lmna* N195K and *Lmna* H222P mice expressing the cGAS-tdTomato transgene. Analysis performed for whole fiber (left) and by classification of nuclei located at the MTJ or within the body of the fiber (right). Data for *Lmna* WT and *Lmna* KO reproduced from Fig. 4E for comparison. *n* = 5-8 mice per genotype, with 5 fibers per animal. ***, *p* < 0.001 *vs*. *Lmna* WT. †, *p* < 0.01 *vs*. nuclei in the muscle body, *, *p* < 0.05 *vs*. *Lmna* WT.

**Supplemental Figure 8. Nuclear envelope rupture in *Lmna* KO muscle fibers is increased at myotendinous junctions**. Representative image of a single isolated muscle fiber demonstrating the enrichment of cGAS+ nuclei at the myotendinous junctions. Scale bar: 200 µm.

**Supplemental Figure 9.** ***Lmna* mutant myonuclei have increased presence of Hsp90 *in vitro* and *in vivo***. (**a**) Representative image of nuclear localization of a large cytosolic protein, Hsp90, inside *Lmna* KO nuclei in myofiber differentiated for 10 days. White arrow indicates a nucleus with no observable chromatin defect and little Hsp90 nuclear accumulation, while the yellow arrow marks a nucleus with a chromatin protrusion and increased nuclear Hsp90 accumulation. Scale bar: 10 µm. (**b**) Quantification of the fluorescence intensity of nuclear Hsp90 levels for *Lmna* WT, *Lmna* KO, *Lmna* KO + Lamin A, *Lmna* N195K and *Lmna* H222P myofibers *in vitro*. For each nucleus, the nuclear fluorescence intensity was normalized to the cytosolic intensity immediately adjacent to each nucleus. *n* = 25-56 nuclei per genotype from 3 independent experiments. ***, *p* < 0.001 *vs*. *Lmna* WT (*p* < 0.001). (**c**) Representative image of Hsp90 nuclear localization in myonuclei from *Lmna* WT and *Lmna* KO mice. Scale bar: 10 µm. (**d**) Quantification of the fluorescence intensity of nuclear HSP90 levels for *Lmna* WT, *Lmna* KO, and *Lmna* H222P isolated single fibers. For each nucleus, the nuclear fluorescence intensity was normalized to the cytosolic intensity immediately beside each nucleus *n* = 25-56 nuclei per genotype from 3 independent experiments. ***, *p* < 0.001 *vs*. *Lmna* WT.

**Supplemental Figure 10. *Lmna* KO MTJ myonuclei have increased DNA-PK activity *in vivo*.** Quantification of p-DNA-PKcs immunofluorescence in isolated muscle fibers from *Lmna* and *Lmna* KO mice. *n* = 4-5 per genotype.

**Supplemental Figure 11. The *Lmna* KO myonuclei with the highest amount of γH2AX foci frequently display chromatin protrusions.** Analysis of DNA damage, assessed by γH2AX staining, in *Lmna* KO nuclei, comparing nuclei with chromatin protrusions to those without protrusions. Chromatin protrusions were assessed based on the presence of chromatin extending beyond the nuclear envelope, marked by lamin B-staining, *n* = 3**.** **, *p* < 0.01 *vs*. no protrusion.

**Supplemental Figure 12. *Lmna* KO myotubes have no defects in DNA damage repair.** (**a**) Representative images of γH2AX foci in *Lmna* WT and *Lmna* KO myotubes at 3, 6 and 24 hours following a 5 Gy dose with radiation or no irradiation control. (**b**) Quantification of γH2AX after 3, 6 and 24 hours post-irradiation or no irradiation control. *n* = 3 independent cell lines. ***, *p* < 0.001 *vs*. control.

**Supplemental Figure 13. Inducing DNA damage or inhibiting DNA repair does not promote additional cells death in *Lmna* KO myofibers.** Quantification of cellular viability in *Lmna* KO myofibers using MTT assay following DNA damage induction with phleomycin, with and without concurrent treatment with DNA-PKi (NU7441) and/or ATMi (KU55933). *n* = 3 independent experiments per condition.

**Supplemental Figure 14.** **Microtubules form cage-like structures around myonuclei**. Representative immunofluorescence image of an isolated *Lmna* WT muscle fiber stained for tubulin (magenta), F-actin (green), and DNA (blue), showing characteristic ‘microtubule cage’ around myonucleus. Scale bar: 5 µm.

**Supplemental Figure 15.** **Inhibiting myofiber contractility does not prevent nuclear envelope rupture in *Lmna* KO myofibers**. (**a**) Quantification of cGAS-mCherry foci formation during 10 day myofiber differentiation follow treatment with nifedipine (5 µM), which inhibits contractility, or DMSO vehicle control, starting at day 5 of differentiation. *N* = 3 independent experiments. (**b**) Quantification of chromatin protrusions at day 7 of differentiation following treatment with nifedipine (5 µM) or DMSO, starting at day 4. Data generated from *n* = 3 independent experiments in which 27-53 nuclei were analyzed per genotype.

**Supplemental Figure 16. Fraction of nuclei with severe chromatin protrusions increased over time in *Lmna* mutant myofibers**. Quantification of the relative distribution of chromatin protrusion lengths in *Lmna* KO, *Lmna* N195K and *Lmna* H222P muscle cells at day 5 and day 10 of *in vitro* differentiation.

**Supplemental Figure 17.** **Kif5b depletion in myotubes reduced chromatin protrusions and DNA damage in *Lmna* KO myonuclei**. (**a**) Western blot for Kif5b in myoblasts treated with a non-target control siRNA (siRNA NT) or siRNA against Kif5b. *n* = 3 independent experiments. (**Bottom**) Corresponding quantification. ***, *p* < 0.001 *vs*. respective genotype siRNA NT control. (**b**) Representative images of *Lmna* KO myofiber at day 5 of differentiation treated with either a non-target control siRNA (siRNA NT) or siRNA against kinesin-1 (siRNA Kif5b) at day 0. Scale bar: 20 µm. Quantification of the number of chromatin protrusions at day 5 of differentiation in *Lmna* KO cells treated with non-target (NT) siRNA or depleted for Kif5b using two independent siRNAs (Kif5b#3 and Kif5b#4). *n* = 4 independent experiments, with 155-270 nuclei counted per image. ***, *p* < 0.001 *vs*. NT control. (**c**) Representative images of *Lmna* KO cells treated with either non-target (NT) siRNA or siRNA against Kif5b and immunofluorescently labeled for γH2AX, showing fewer chromatin protrusions and less DNA damage in the Kif5b depleted cells. Scale bar: 20 µm (**d**) Quantification of the number of γH2AX foci in *Lmna* KO myonuclei following treatment with either non-target siRNA or siRNA against Kif5b. *n* = 3 independent experiments in which 27-53 nuclei are counted per image.

**Supplemental Figure 18. Expression of the DN-KASH2 construct disrupts the LINC complex and limits nuclear movement, without affecting myofiber function in *Lmna* WT myofibers.** (**a**) Representative image showing displacement of endogenous nesprin-1 in myofibers expressing the DN-KASH2 construct, and no displacement of nesprin-1 in myofibers expressing the DN-KASH2ext construct. Scale bar: 10 µm. (**b**) Representative image showing nuclear clustering in myofibers expressing the DN-KASH2 construct, and normal nuclear spreading in myofibers expressing the DN-KASH2ext construct. Scale bar: 20 µm. (**c**) Quantification of cell viability following DN-KASH2 or DN-KASH2ext treatment in *Lmna* WT cells using the MTT assay. *n* = 6 per condition from 3 independent experiments. (**d**) Quantification of myofiber contraction following DN-KASH2 or DN-KASH2ext treatment in *Lmna* WT cells based on the percent of contractile fibers. *n* = 4 independent experiments per condition. (**e**) Quantification of the number of chromatin protrusions in *Lmna* KO myonuclei expressing either DN-KASH2 or DN-KASH2ext. *n* = 3 per condition. ***, *p* < 0.001 vs. DN-KASH2 + DOX; †††, *p* < 0.001 vs DN-KASH2ext ‒ DOX and DN-KASH2ext + DOX. **, *p* < 0.01 vs. DN-KASH2ext + DOX. (**f**) Quantification of the extent of DNA damage based on the number of γH2AX foci per nucleus during myofiber differentiation. *Lmna* KO myonuclei expressing the DN-KASH2 construct show a decrease in the nuclei with >25 foci. *n* = 4 per condition.

**Supplemental Figure 19.** **Human laminopathy muscle tissue shows increased 53BP1 staining.** Representative images of cryopreserved human muscle biopsy tissue from individuals with *LMNA* muscular dystrophy and age-matched controls stained with either (**a**) anti-53BP1, DAPI, and Phalloidin, or (**b**) anti-53BP1, anti-dystrophin, and DAPI. The boxed regions in the left column of (b) are magnified in the right column, showing increased anti-53BP1 staining in muscle from laminopathy individuals versus age-matched controls.

**Supplemental Figure 20. Proposed mechanism by which *Lmna* mutations result in myofiber dysfunction and death.** (**a**) Kinesin-1 motor proteins spread myonuclei along the myotubes axis during differentiation. In *Lmna* mutant cells, which have mechanically weaker nuclei, the localized forces associated with nuclear migration cause chromatin protrusion and NE ruptures. This mechanically induced nuclear damage results in DNA damage, detected by H2AX foci, and activation of the DNA damage response pathways, which leads to decline in myofiber health and cell death. (**b**) Schematic flow chart delineating the steps described in panel A, along with interventions explored in this work. Stabilizing microtubules surrounding the myonuclei reinforces the *Lmna* mutant nuclei and prevents chromatin protrusions and NE ruptures. Inhibiting nuclear movement by Kif5b depletions similarly prevents nuclear damage. Muscle contractions may also contribute to nuclear damage *in vivo*.

**Supplemental Movies**

**Supplemental Movies 1-4.** Representative movies of spontaneous contractions in *Lmna* WT, *Lmna* KO, *Lmna* N195K, and *Lmna* H222P myofibers after 10 days of differentiation.

**Supplemental Movie 5.** Representative movie of micropipette aspiration of *Lmna* WT, *Lmna* KO *Lmna* N195K, and *Lmna* H222P myoblasts.

**Supplemental Movie 6.** Representative movie of microharpoon manipulation of *Lmna* WT and *Lmna* KO myotubes after day 5 of differentiation.

**Supplemental Movie 7.** Time-lapse of nuclear envelope rupture in *Lmna* KO myotubes after four days of differentiation. Note the loss of soluble NLS-GFP from the nucleus into the cytoplasm.

**Supplemental Movie 8.** Representative movie of microharpoon manipulation of *Lmna* KO myotubes after day 5 of differentiation following 24 hours of treatment with either 50 nM paclitaxel or DMSO control.

**Supplemental Movie 9.** Time-lapse of nuclear envelope rupture during myonuclear movement at 5 days of differentiation. Note the loss of NLS-GFP from the nucleus is immediately followed by the formation of cGAS-mCherry foci at the site of rupture.

**Supplemental Movies 10-13.** Representative movies of spontaneous contractions in *Lmna* WT myofibers after 10 days of differentiation expressing a doxycycline inducible GFP-KASH2 to disrupt nucleo-cytoskeletal force transmission or the GFP-KASH2ext control, and the corresponding non-doxycycline induced controls.

**Supplemental Movies 14-17.** Representative movies of spontaneous contractions in *Lmna KO* myofibers after 10 days of differentiation expressing a doxycycline inducible GFP-KASH2 to disrupt nucleo-cytoskeletal force transmission or the GFP-KASH2ext control, and the corresponding non-doxycycline induced controls.

**Supplemental Table**

| Antibody | Cat# | Vendor | Dilution |
| --- | --- | --- | --- |
| MyHC | A4.1025 | DSHB | 1:100 |
| MyHC | MAB4470-SP | Novus Biologicals | 1:500 |
| Lamin B (M-20) | sc-6217 | Santa Cruz | 1:200 |
| Lamin B1 (B-10) | sc-374015 | Santa Cruz | 1:200 |
| Lamin A (H-102) | sc-20680 | Santa Cruz | 1:200 |
| Lamin A/C (E1) | sc-376248 | Santa Cruz | 1:200 |
| Gamma-H2AX (Ser139) | 80312 | Cell Signaling | 1:200 |
| Gamma-H2AX (Ser139) | 9718 | Cell Signaling | 1:200 |
| HSP90 α/β (F-8) | sc-13119 | Santa Cruz | 1:200 |
| Nesprin1-E | MANNES1E | Glen Morris | 1:500 |
| Nesprin-1-A | MANNES1A | Glen Morris | 1:500 |
| alpha-tubulin | T9026 | Sigma | 1:500(IF) 1:5000(WB) |
| NPC (414) | Ab50008 | Abcam | 1:500 |
| Emerin | NCL-EMERIN | Leica | 1:200 |
| DNA-PKcs (S2056) | ab18192 | Abcam | 1:1000 |
| DNA-PKcs | sc-390849 | Santa Cruz | 1:750 |
| Cleaved Caspase-3 | 9661 | Cell Signaling | 1:500 |
| 53BP1 | NB100-304 | Novus Biologicals | 1:1000 |
| Dystrophin | Mab7A10 | University of Iowa Hospitals and Clinics Pathology Core | 1:20 |

**Supplemental Table 1. Antibodies and corresponding dilutions.** Primary antibodies for immunofluorescence staining and western blotting.
