## Supplementary Figures for "Mutant lamins cause nuclear envelope rupture and DNA damage in skeletal muscle cells"

### Supp Fig. 1

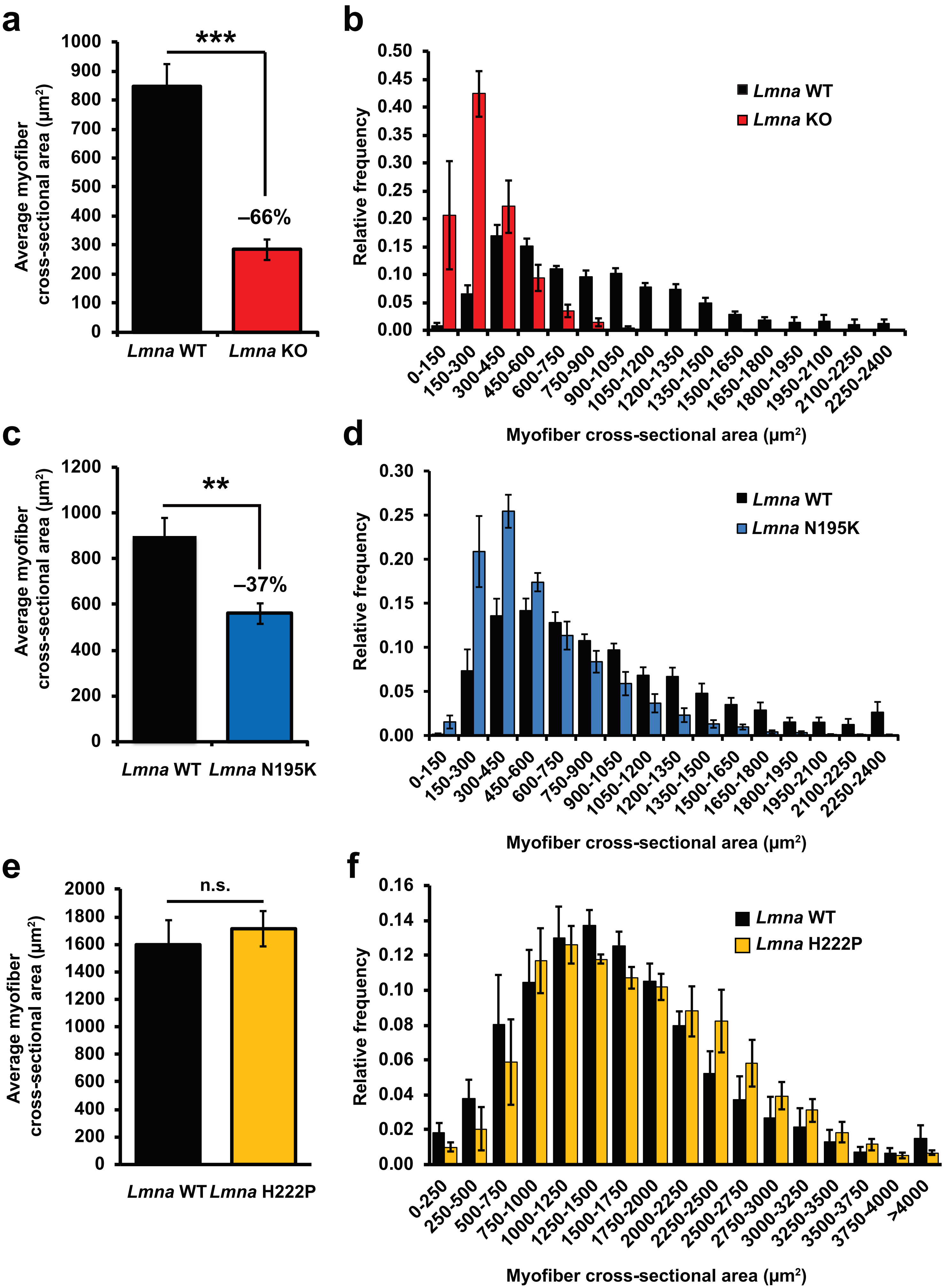

a
Average myofiber cross-sectional area (µm²)
1000
900
800
700
600
500
400
300
200
100
0
***
–66%
Lmna WT
Lmna KO
b
Relative frequency
0.50
0.45
0.40
0.35
0.30
0.25
0.20
0.15
0.10
0.05
0.00
Lmna WT
Lmna KO
0-150
150-300
300-450
450-600
600-750
750-900
900-1050
1050-1200
1200-1350
1350-1500
1500-1650
1650-1800
1800-1950
1950-2100
2100-2250
2250-2400
Myofiber cross-sectional area (µm²)
c
Average myofiber cross-sectional area (µm²)
1200
1000
800
600
400
200
0
**
–37%
Lmna WT
Lmna N195K
d
Relative frequency
0.30
0.25
0.20
0.15
0.10
0.05
0.00
Lmna WT
Lmna N195K
0-150
150-300
300-450
450-600
600-750
750-900
900-1050
1050-1200
1200-1350
1350-1500
1500-1650
1650-1800
1800-1950
1950-2100
2100-2250
2250-2400
Myofiber cross-sectional area (µm²)
e
Average myofiber cross-sectional area (µm²)
2000
1800
1600
1400
1200
1000
800
600
400
200
0
n.s.
Lmna WT
Lmna H222P
f
Relative frequency
0.16
0.14
0.12
0.10
0.08
0.06
0.04
0.02
0.00
Lmna WT
Lmna H222P
0-250
250-500
500-750
750-1000
1000-1250
1250-1500
1500-1750
1750-2000
2000-2250
2250-2500
2500-2750
2750-3000
3000-3250
3250-3500
3500-3750
3750-4000
>4000
Myofiber cross-sectional area (µm²)

### Supp Fig. 2

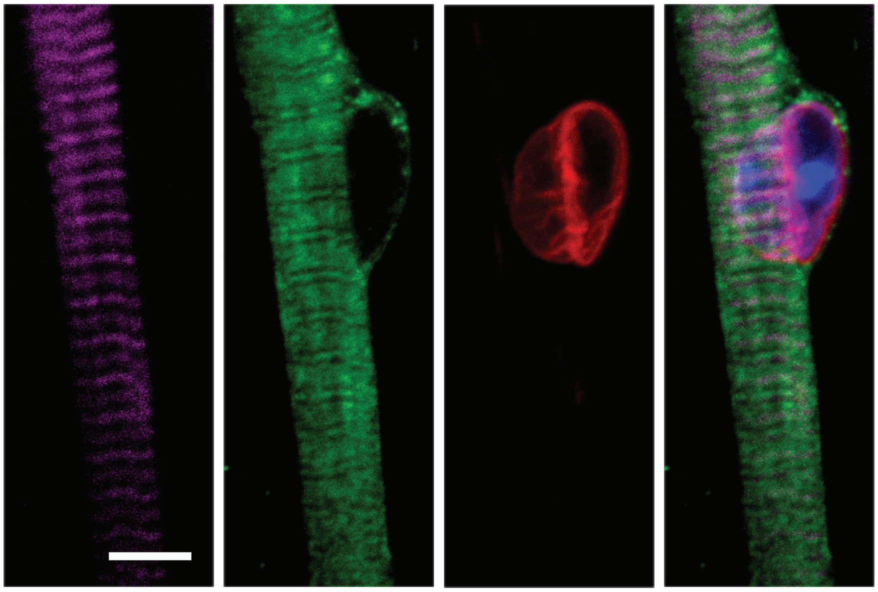

Actin  Myosin heavy chain  Lamin B1  DNA

### Supp Fig. 3

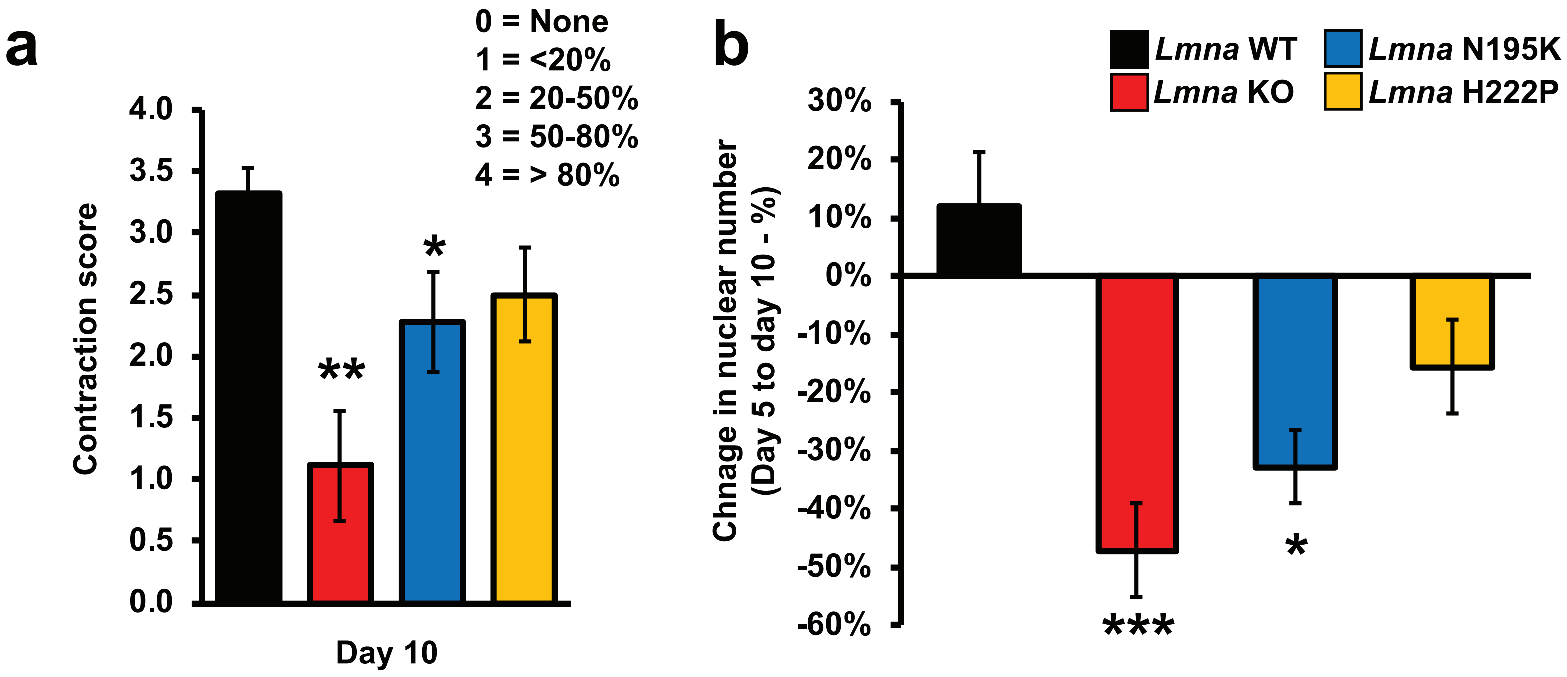

a
0 = None
1 = <20%
2 = 20-50%
3 = 50-80%
4 = > 80%
Contraction score
4.0
3.5
3.0
2.5
2.0
1.5
1.0
0.5
0.0
**
*
Day 10
b
Lmna WT
Lmna KO
Lmna N195K
Lmna H222P
Chnage in nuclear number
(Day 5 to day 10 - %)
30%
20%
10%
0%
-10%
-20%
-30%
-40%
-50%
-60%
***
*

### Supp Fig. 4

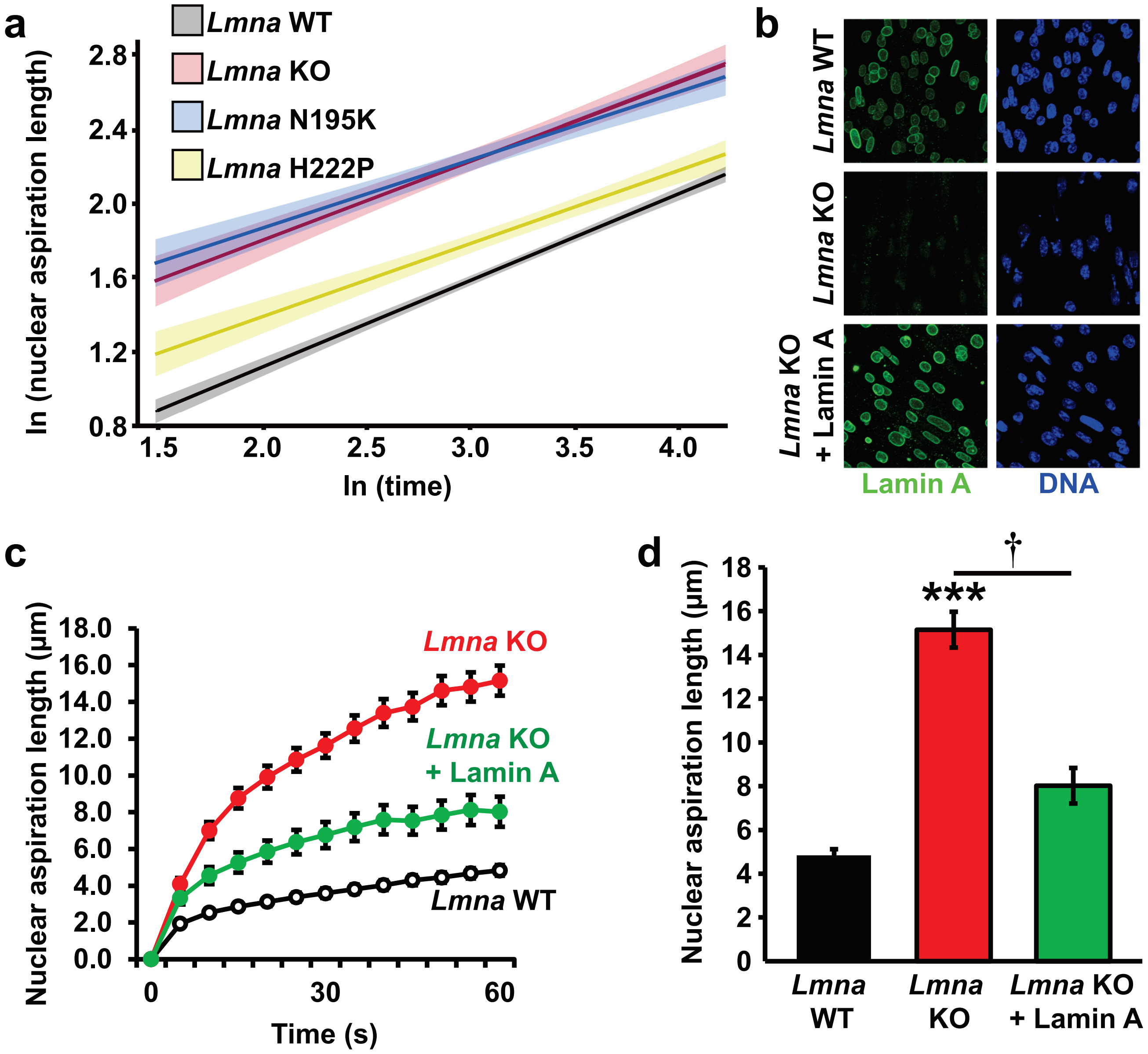
a
Lmna WT
Lmna KO
Lmna N195K
Lmna H222P
ln (nuclear aspiration length)
2.8
2.4
2.0
1.6
1.2
0.8
1.5
2.0
2.5
3.0
3.5
4.0
ln (time)
b
Lmna WT
Lmna KO
Lmna KO + Lamin A
Lamin A
DNA
c
Nuclear aspiration length (µm)
18.0
16.0
14.0
12.0
10.0
8.0
6.0
4.0
2.0
0.0
0
30
60
Time (s)
Lmna KO
Lmna KO + Lamin A
Lmna WT
d
†
***
Nuclear aspiration length (µm)
18
16
14
12
10
8
6
4
2
0
Lmna WT
Lmna KO
Lmna KO + Lamin A

### Supp Fig. 5

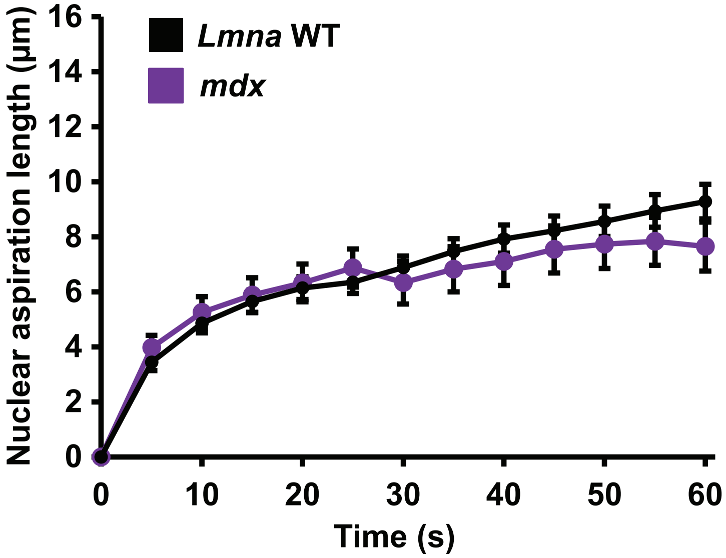

Lmna WT
mdx
Nuclear aspiration length (µm)
16
14
12
10
8
6
4
2
0
0
10
20
30
40
50
60
Time (s)

### Supp Fig. 6

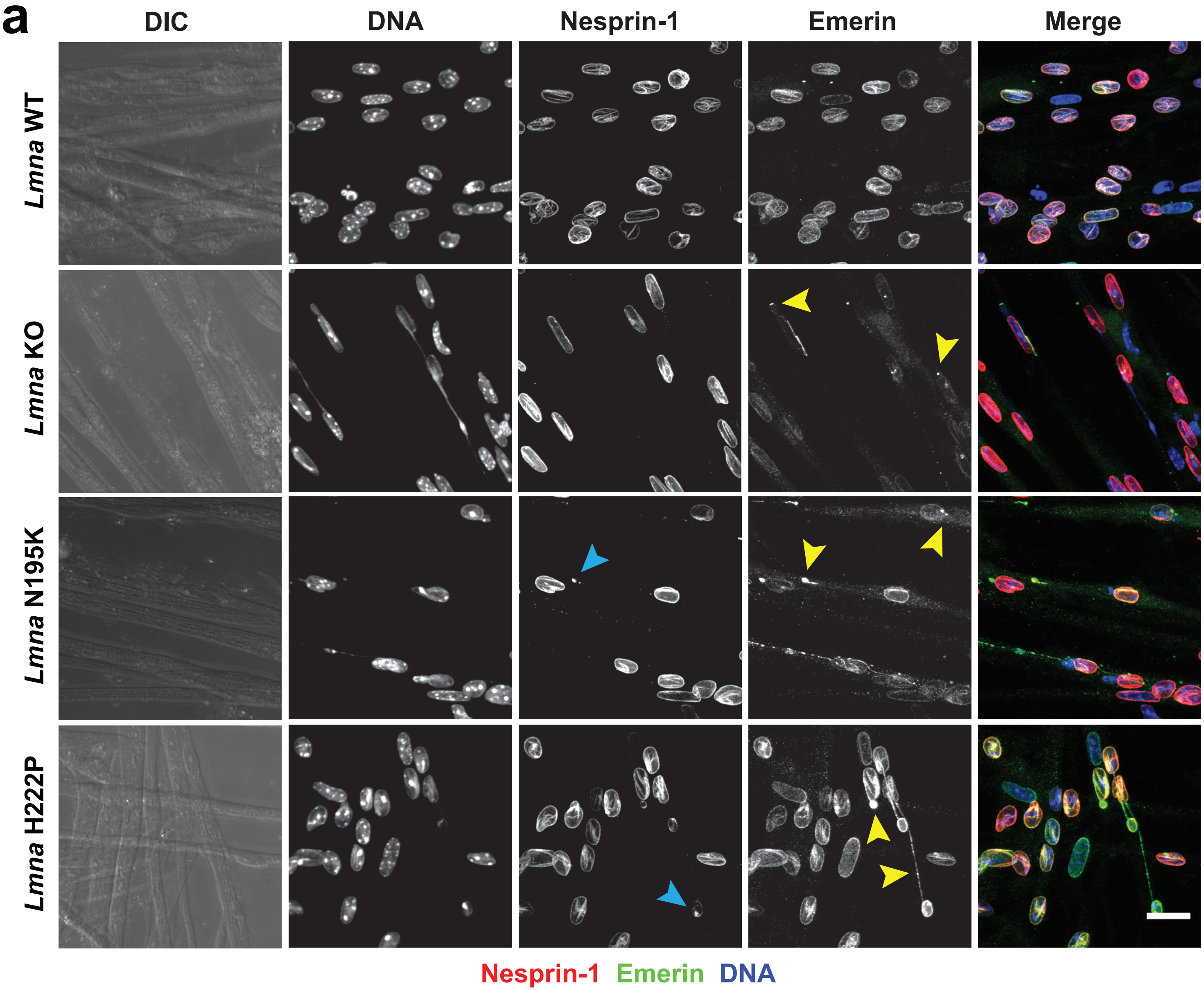
a
DIC
DNA
Nesprin-1
Emerin
Merge
Lmna WT
Lmna KO
Lmna N195K
Lmna H222P
Nesprin-1 Emerin DNA

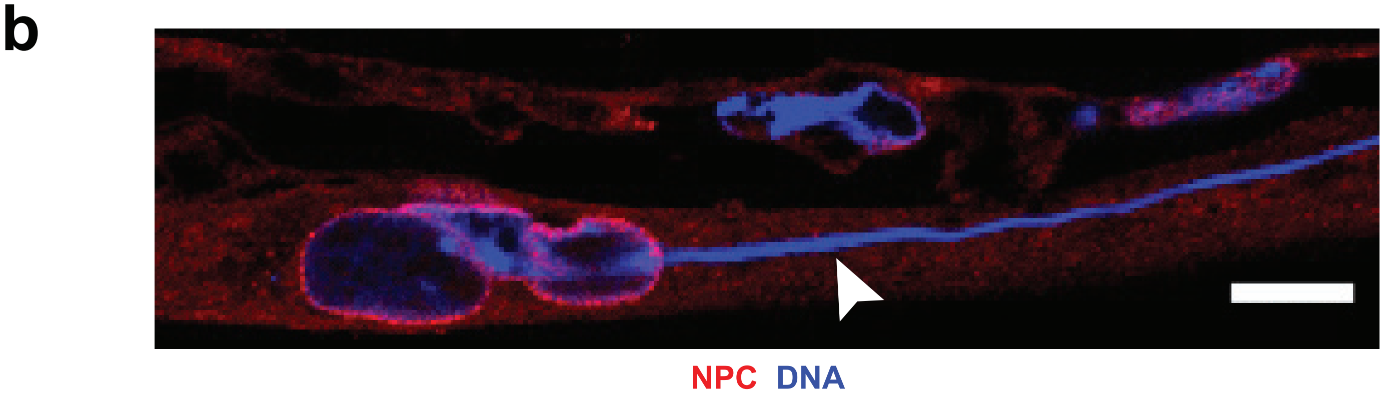
b
NPC DNA

### Supp Fig. 7

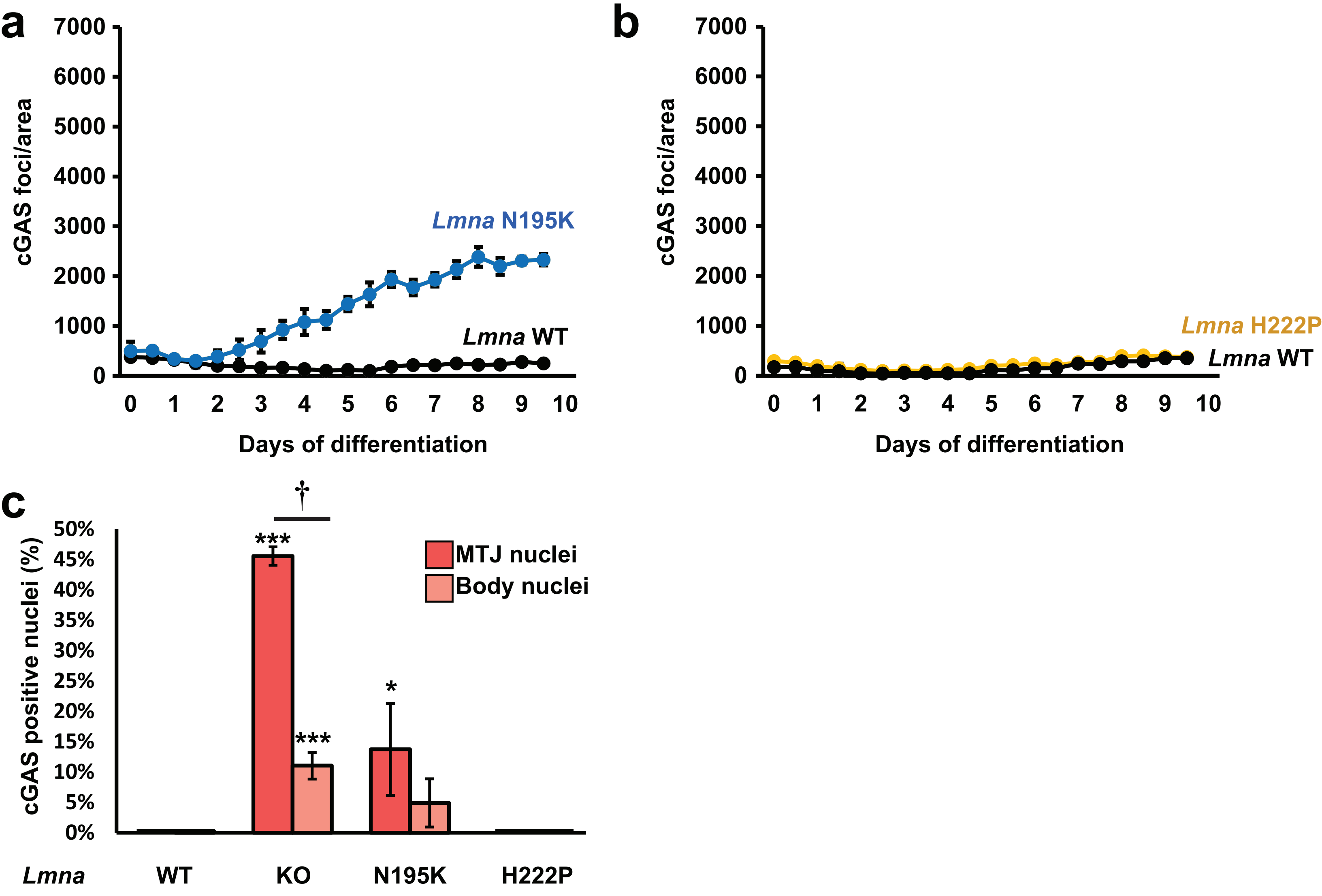

a
cGAS foci/area
7000
6000
5000
4000
3000
2000
1000
0
*Lmna* N195K
*Lmna* WT
0 1 2 3 4 5 6 7 8 9 10
Days of differentiation
b
cGAS foci/area
7000
6000
5000
4000
3000
2000
1000
0
*Lmna* H222P
*Lmna* WT
0 1 2 3 4 5 6 7 8 9 10
Days of differentiation
c
†
***
***
*
cGAS positive nuclei (%)
50%
45%
40%
35%
30%
25%
20%
15%
10%
5%
0%
MTJ nuclei
Body nuclei
*Lmna*
WT
KO
N195K
H222P

**Supp Fig. 8**

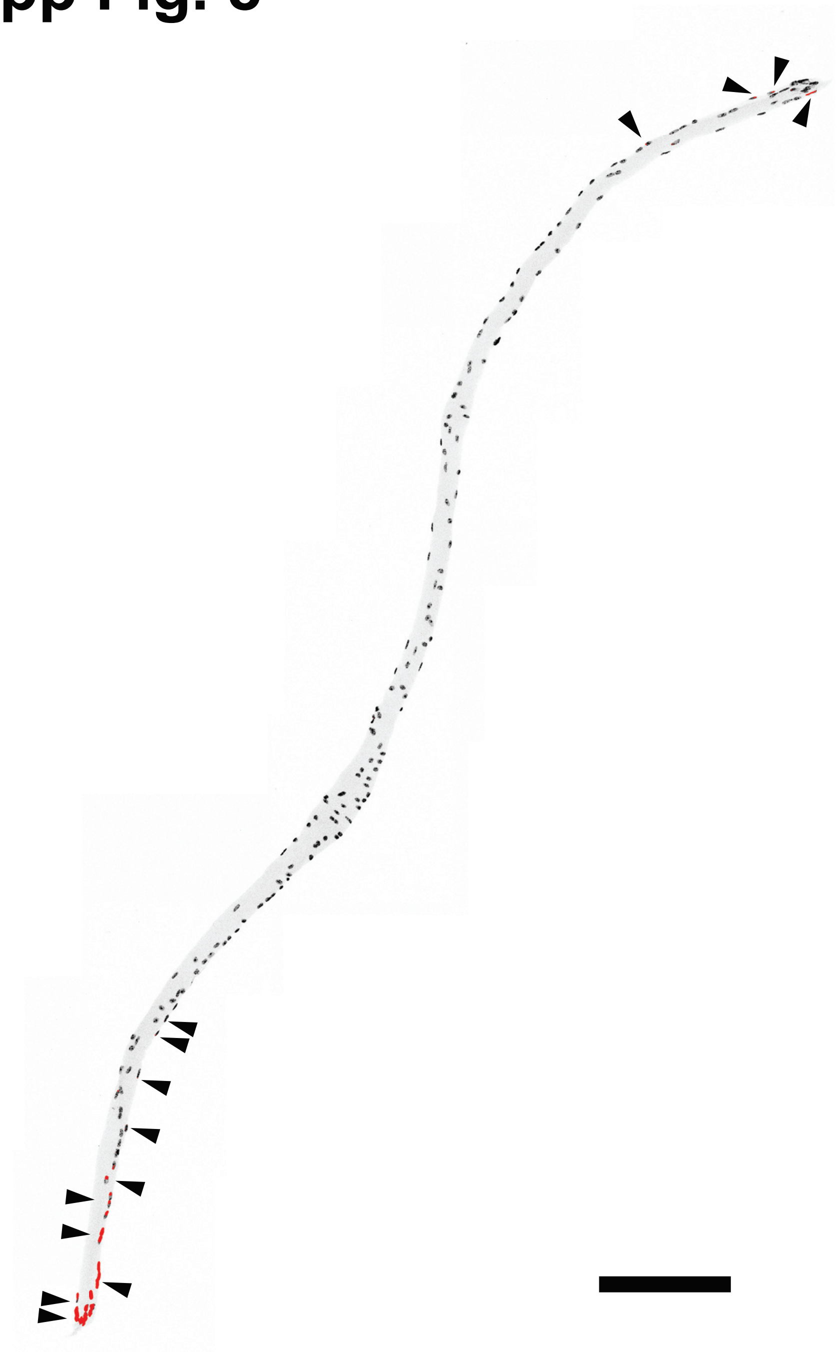

### Supp Fig. 9

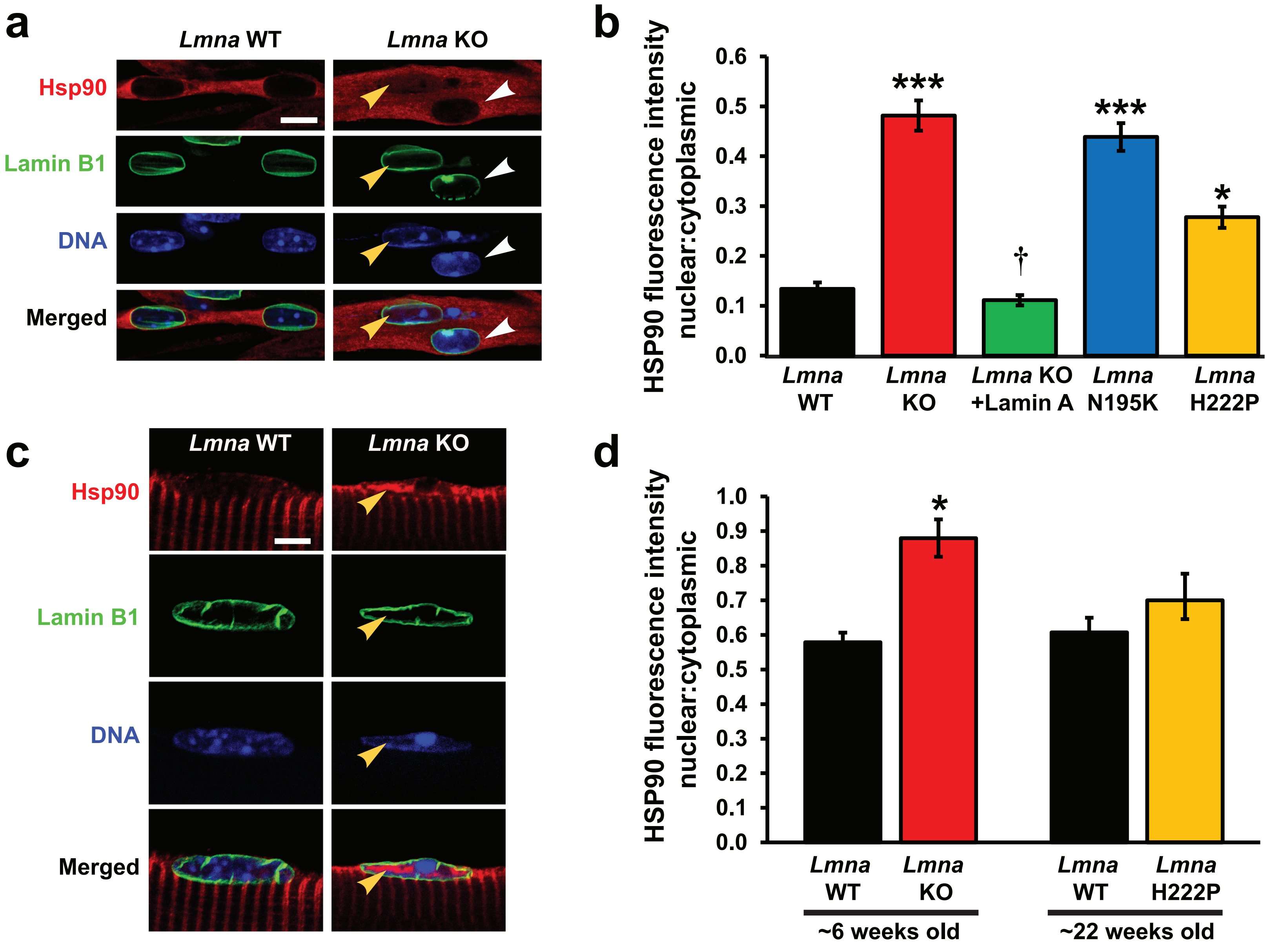
a
Lmna WT
Lmna KO
Hsp90
Lamin B1
DNA
Merged
b
HSP90 fluorescence intensity nuclear:cytoplasmic
0.6
0.5
0.4
0.3
0.2
0.1
0.0
***
***
†
*
Lmna WT
Lmna KO
Lmna KO +Lamin A
Lmna N195K
Lmna H222P
c
Lmna WT
Lmna KO
Hsp90
Lamin B1
DNA
Merged
d
HSP90 fluorescence intensity nuclear:cytoplasmic
1.0
0.9
0.8
0.7
0.6
0.5
0.4
0.3
0.2
0.1
0.0
*
Lmna WT
Lmna KO
Lmna WT
Lmna H222P
~6 weeks old
~22 weeks old

### Supp Fig. 10

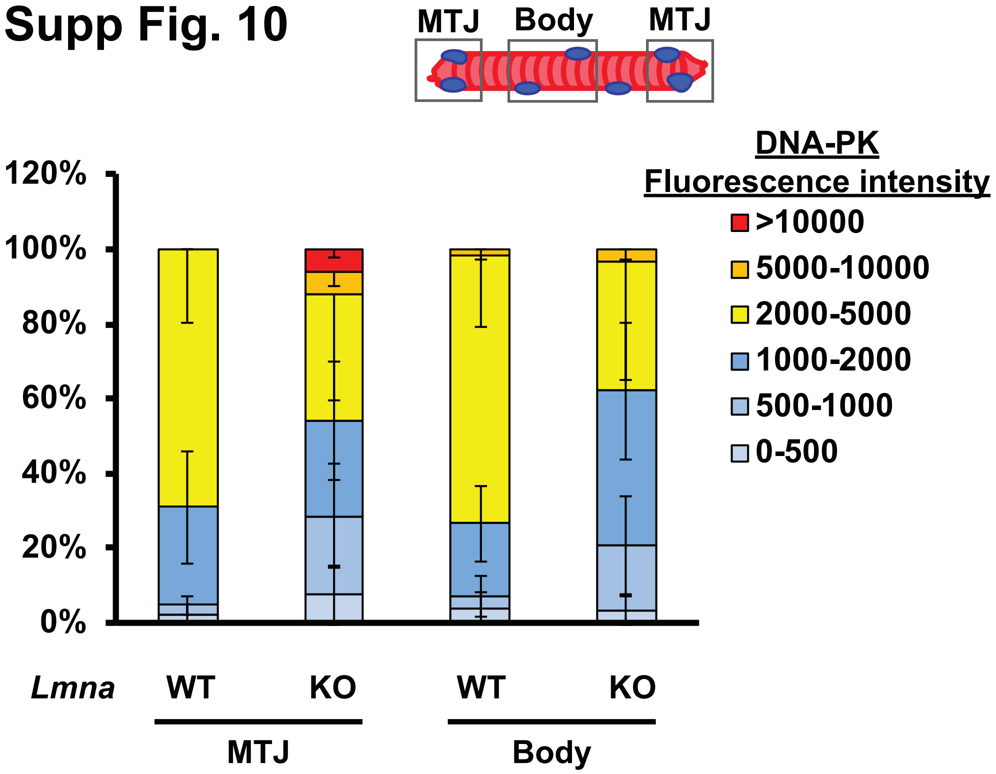
MTJ
Body
MTJ
DNA-PK
Fluorescence intensity
>10000
5000-10000
2000-5000
1000-2000
500-1000
0-500
120%
100%
80%
60%
40%
20%
0%
Lmna
WT
KO
WT
KO
MTJ
Body

### Supp Fig. 11

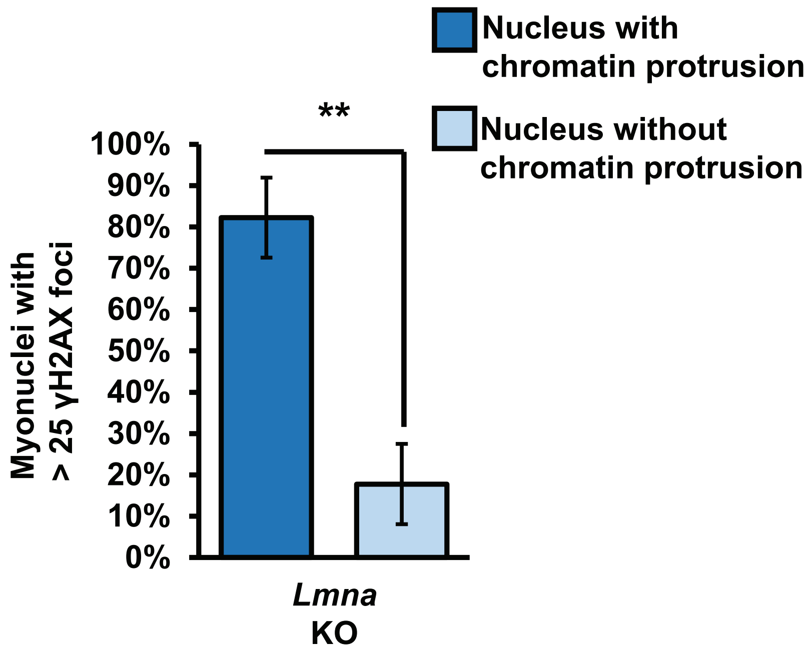

Nucleus with chromatin protrusion
Nucleus without chromatin protrusion
**
100%
90%
80%
70%
60%
50%
40%
30%
20%
10%
0%
Myonuclei with > 25 γH2AX foci
Lmna KO

### Supp Fig. 12

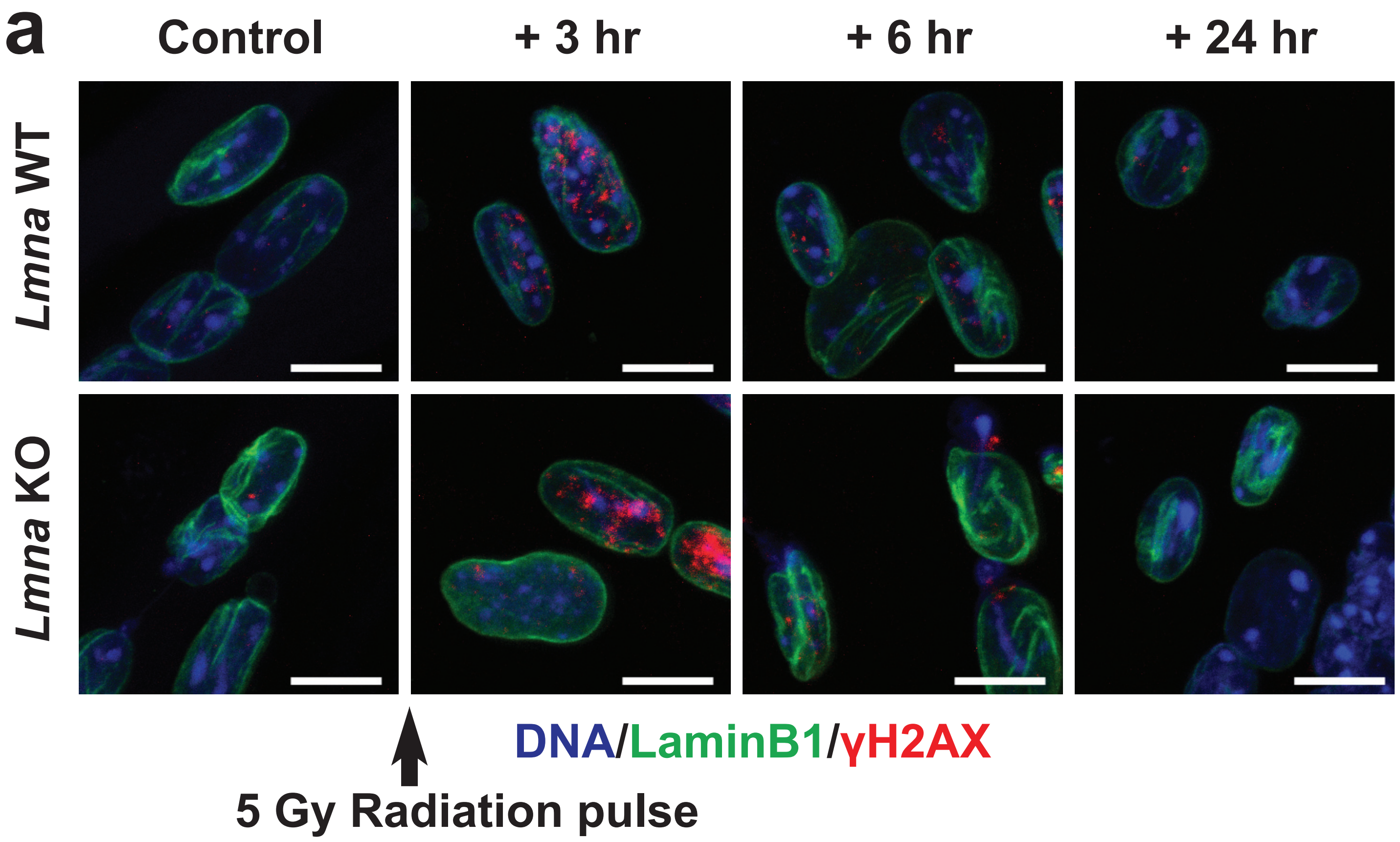
a
Control
+ 3 hr
+ 6 hr
+ 24 hr
Lmna WT
Lmna KO
DNA/LaminB1/γH2AX
5 Gy Radiation pulse

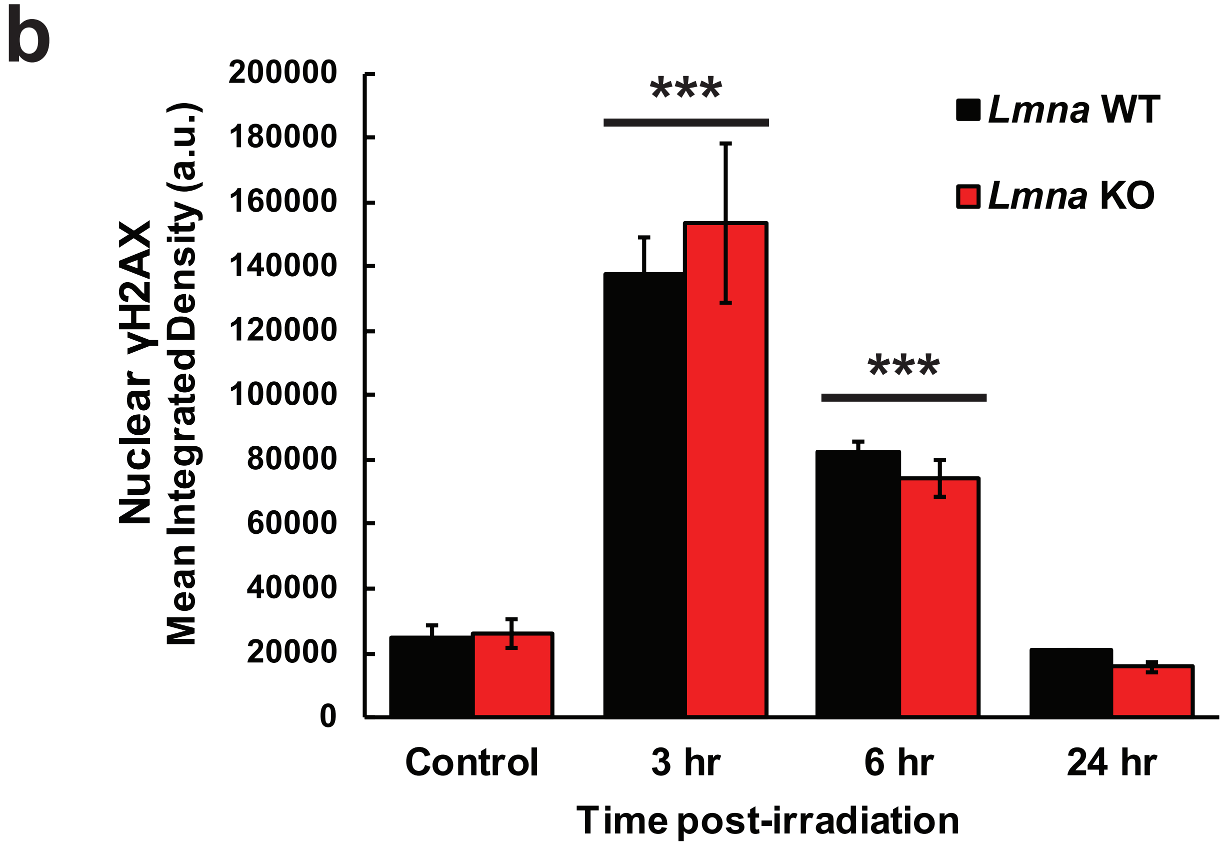
b
Nuclear γH2AX
Mean Integrated Density (a.u.)
200000
180000
160000
140000
120000
100000
80000
60000
40000
20000
0
***
***
Lmna WT
Lmna KO
Control
3 hr
6 hr
24 hr
Time post-irradiation

### Supp Fig. 13

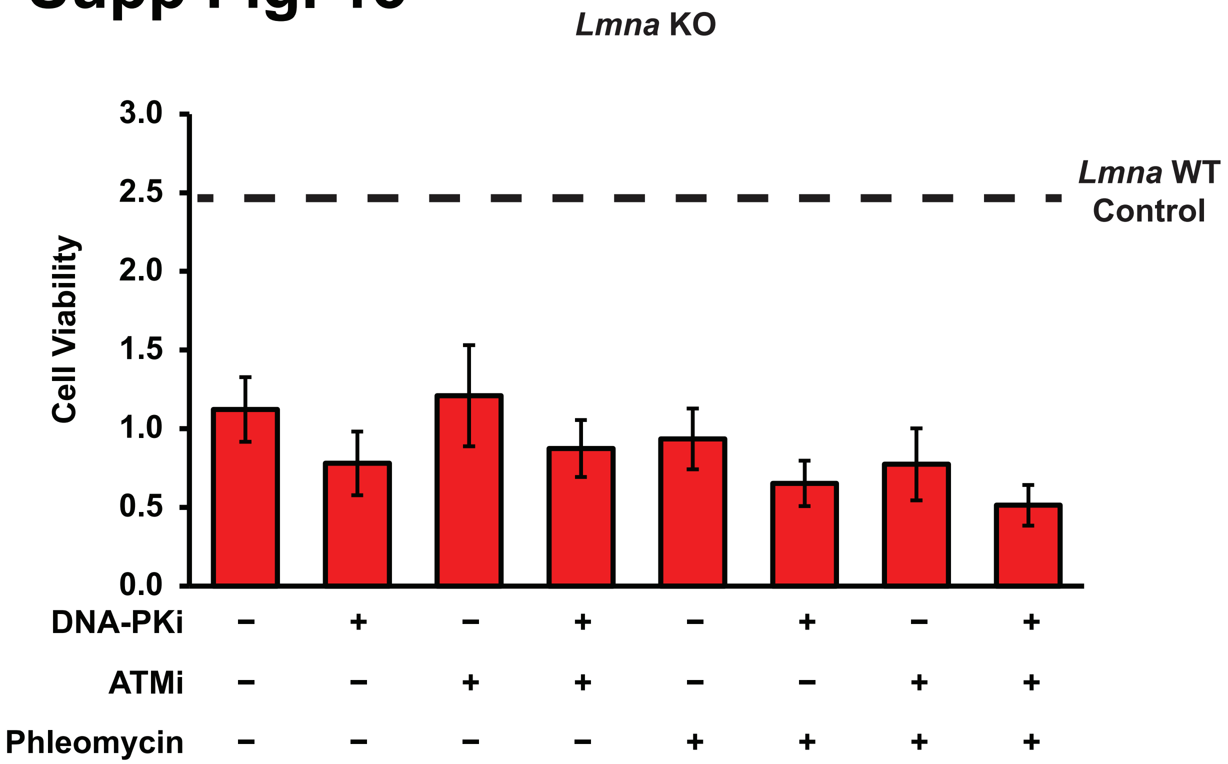
*Lmna* KO
3.0
2.5
2.0
1.5
1.0
0.5
0.0
Cell Viability
*Lmna* WT
Control
DNA-PKi − + − + − + − +
ATMi − − + + − − + +
Phleomycin − − − − + + + +

### Supp Fig. 14

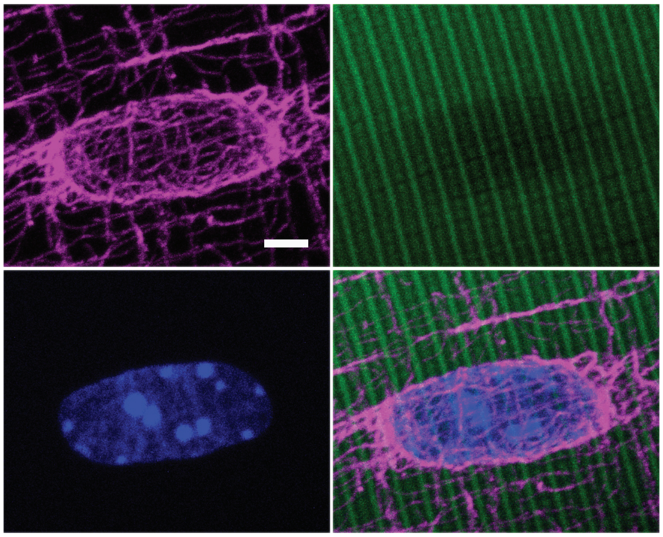

α-Tubulin Actin DNA

### Supp Fig. 15

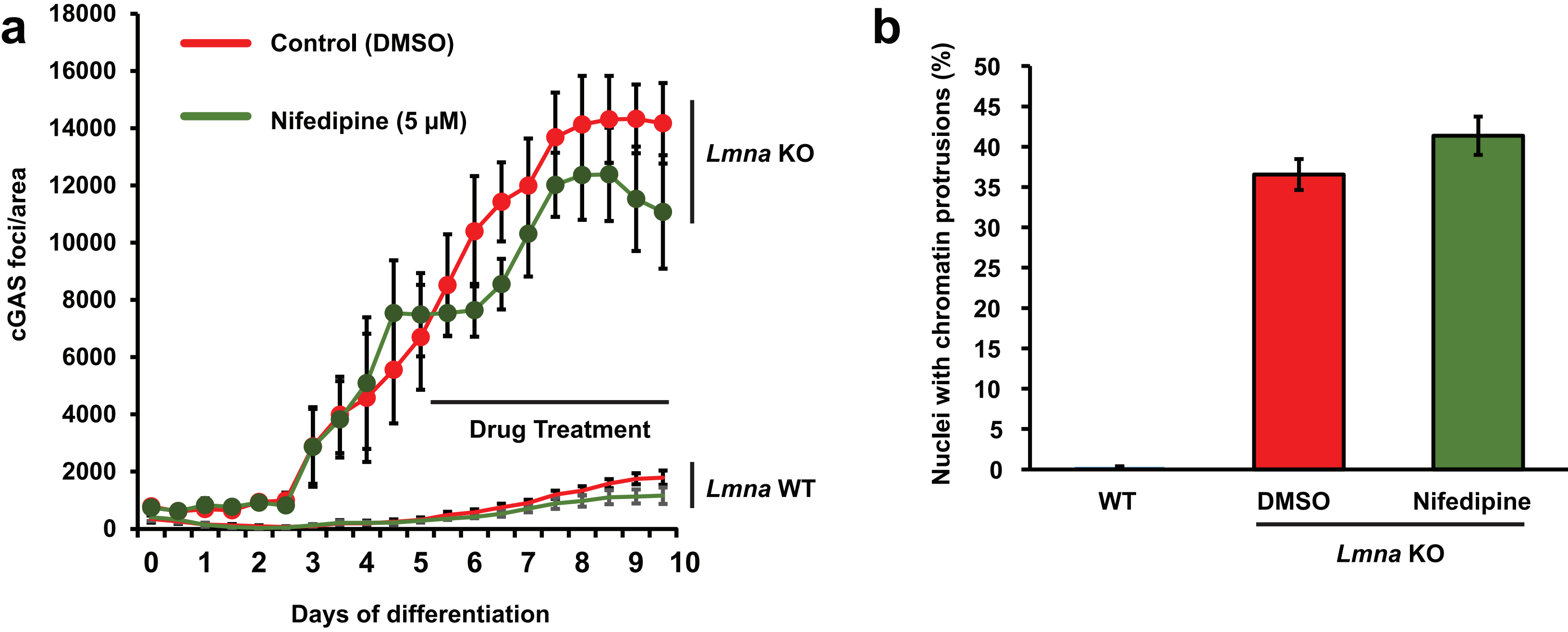
a
Control (DMSO)
Nifedipine (5 μM)
Lmna KO
Lmna WT
Drug Treatment
cGAS foci/area
18000
16000
14000
12000
10000
8000
6000
4000
2000
0
0
1
2
3
4
5
6
7
8
9
10
Days of differentiation
b
Nuclei with chromatin protrusions (%)
50
45
40
35
30
25
20
15
10
5
0
WT
DMSO
Nifedipine
Lmna KO

### Supp Fig. 16

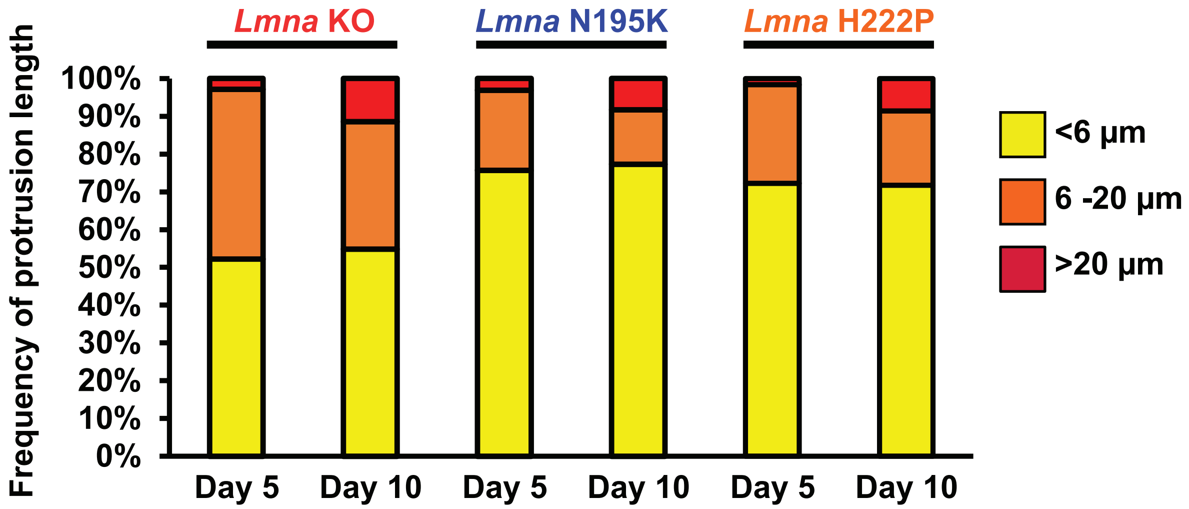

*Lmna* KO
*Lmna* N195K
*Lmna* H222P
Frequency of protrusion length
100%
90%
80%
70%
60%
50%
40%
30%
20%
10%
0%
Day 5
Day 10
Day 5
Day 10
Day 5
Day 10
<6 µm
6 -20 µm
>20 µm

### Supp Fig. 17

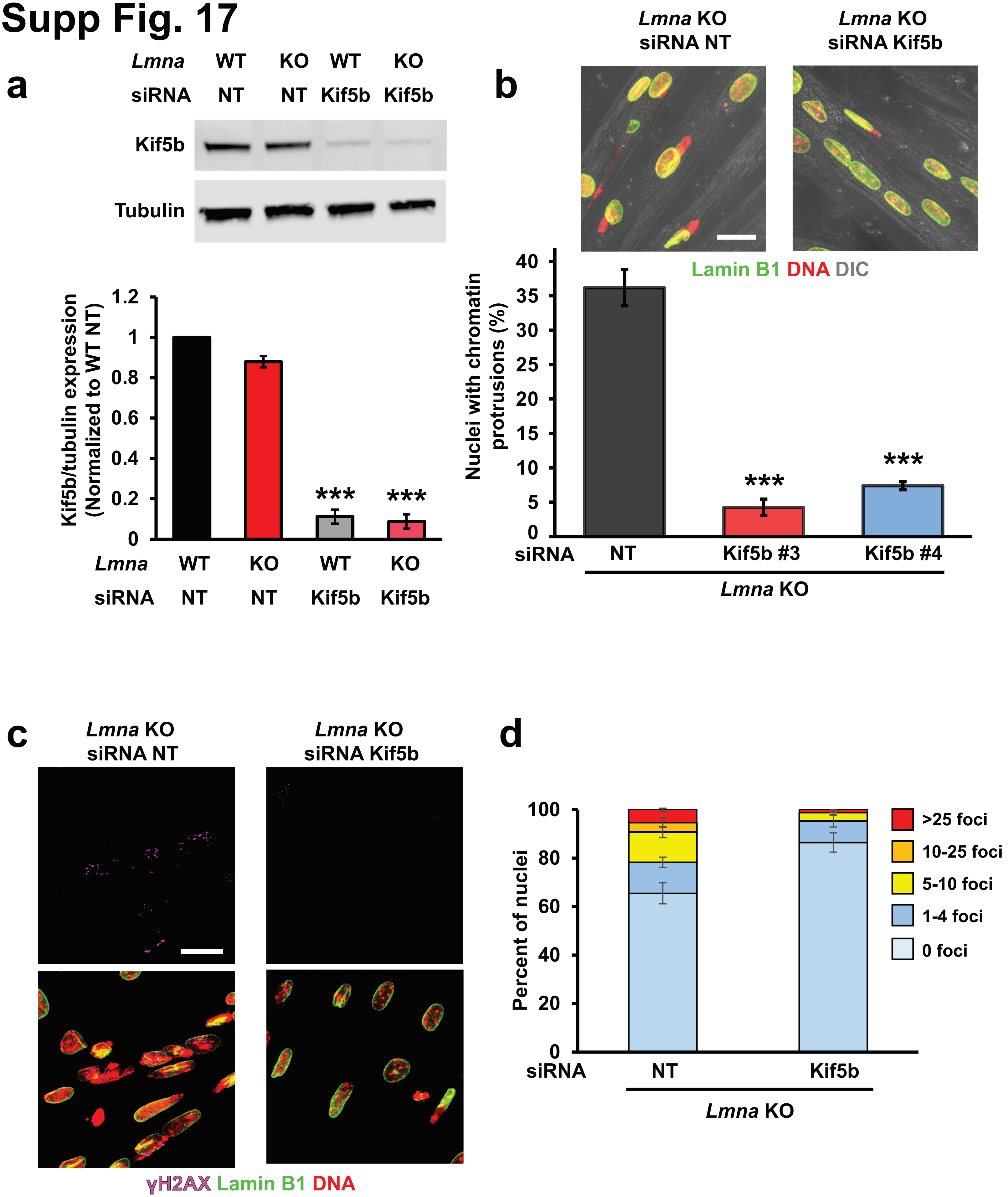
a
Lmna WT KO WT KO
siRNA NT NT Kif5b Kif5b
Kif5b
Tubulin
Kif5b/tubulin expression (Normalized to WT NT)
1.2
1
0.8
0.6
0.4
0.2
0
***
***
Lmna WT KO WT KO
siRNA NT NT Kif5b Kif5b
b
Lmna KO siRNA NT
Lmna KO siRNA Kif5b
Lamin B1 DNA DIC
Nuclei with chromatin protrusions (%)
40
35
30
25
20
15
10
5
0
***
***
siRNA NT Kif5b #3 Kif5b #4
Lmna KO
c
Lmna KO siRNA NT
Lmna KO siRNA Kif5b
γH2AX Lamin B1 DNA
d
Percent of nuclei
100
80
60
40
20
0
>25 foci
10-25 foci
5-10 foci
1-4 foci
0 foci
siRNA NT Kif5b
Lmna KO

### Supp Fig. 18

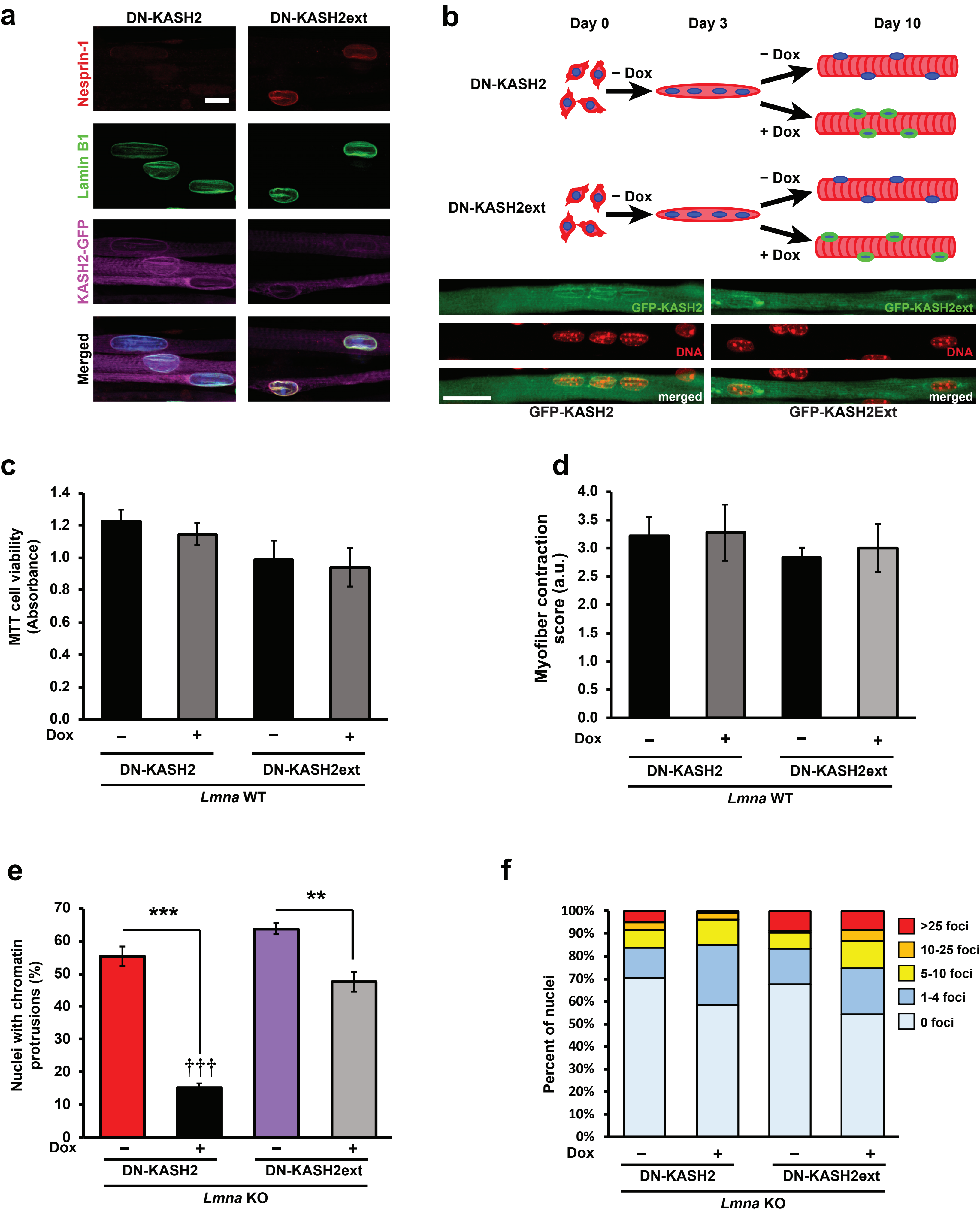

a
DN-KASH2
DN-KASH2ext
Nesprin-1
Lamin B1
KASH2-GFP
Merged
b
Day 0
Day 3
Day 10
DN-KASH2
DN-KASH2ext
− Dox
+ Dox
GFP-KASH2
GFP-KASH2ext
DNA
merged
GFP-KASH2
GFP-KASH2Ext
c
MTT cell viability (Absorbance)
1.4
1.2
1.0
0.8
0.6
0.4
0.2
0.0
Dox
–
+
DN-KASH2
DN-KASH2ext
Lmna WT
d
Myofiber contraction score (a.u.)
4.0
3.5
3.0
2.5
2.0
1.5
1.0
0.5
0.0
Dox
–
+
DN-KASH2
DN-KASH2ext
Lmna WT
e
Nuclei with chromatin protrusions (%)
70
60
50
40
30
20
10
0
***
**
†††
Dox
–
+
DN-KASH2
DN-KASH2ext
Lmna KO
f
Percent of nuclei
100%
90%
80%
70%
60%
50%
40%
30%
20%
10%
0%
>25 foci
10-25 foci
5-10 foci
1-4 foci
0 foci
Dox
–
+
DN-KASH2
DN-KASH2ext
Lmna KO

### Supp Fig. 19

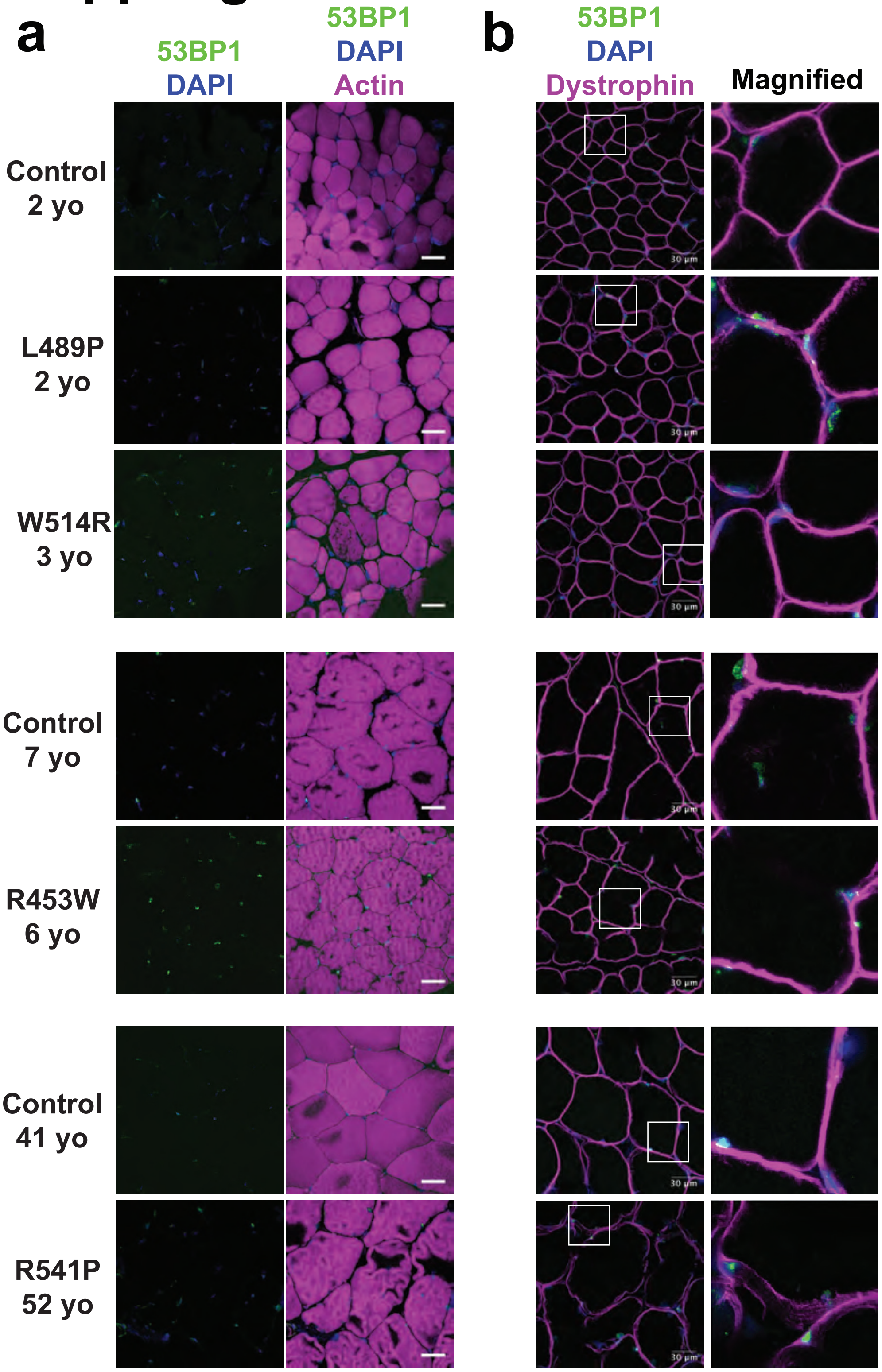

a
b
53BP1
DAPI
53BP1
DAPI
Actin
53BP1
DAPI
Dystrophin
Magnified
Control
2 yo
L489P
2 yo
W514R
3 yo
Control
7 yo
R453W
6 yo
Control
41 yo
R541P
52 yo
30 µm

### Suppl. Fig 20

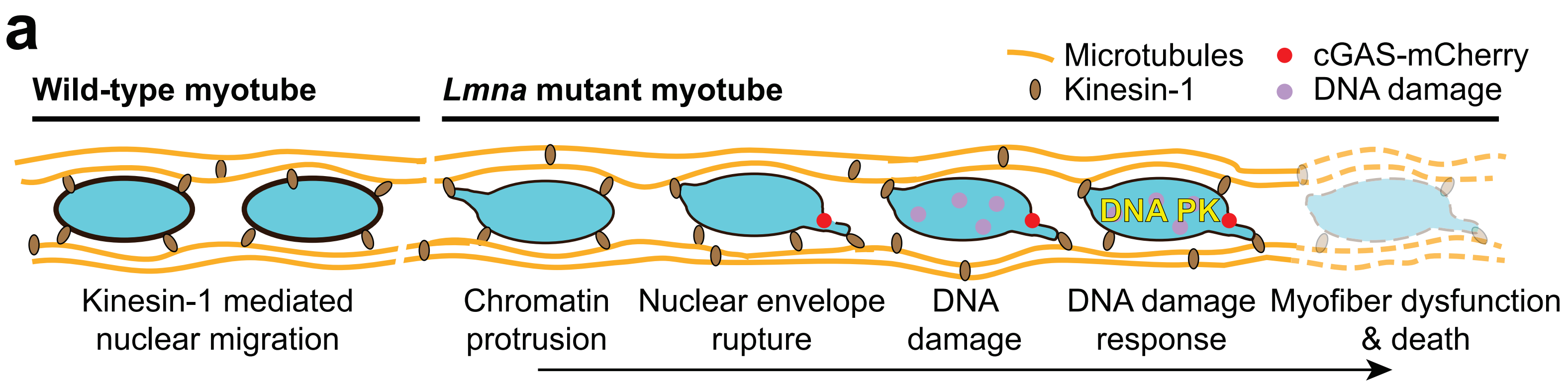
a
Wild-type myotube
*Lmna* mutant myotube
Microtubules
Kinesin-1
cGAS-mCherry
DNA damage
DNA PK
Kinesin-1 mediated nuclear migration
Chromatin protrusion
Nuclear envelope rupture
DNA damage
DNA damage response
Myofiber dysfunction & death

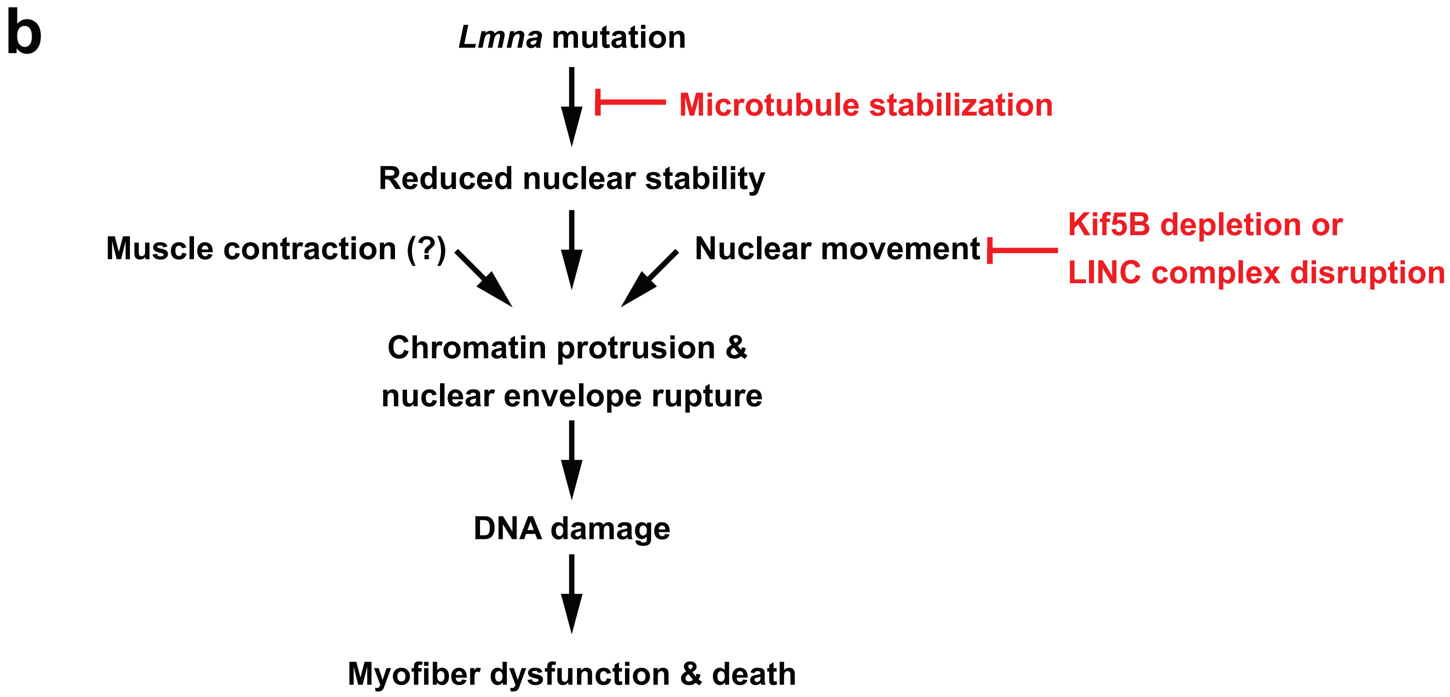
b
*Lmna* mutation
Microtubule stabilization
Reduced nuclear stability
Muscle contraction (?)
Nuclear movement
Kif5B depletion or
LINC complex disruption
Chromatin protrusion &
nuclear envelope rupture
DNA damage
Myofiber dysfunction & death
